# Durability and diversity of human upper airway T cell memory

**DOI:** 10.64898/2026.09.15.751278

**Authors:** Sydney I. Ramirez, Farhoud Faraji, Paul G. Lopez, Lucas Garin-Ortega, Natalie Hern, Osirus Eisenman, Ahmed Azhan, April Frazier, Elizabeth J. Phillips, Simon Mallal, Alessandro Sette, Thamotharampillai Dileepan, Marc K. Jenkins, Shane Crotty

## Abstract

T cells are major contributors to protective immunity against pathogens and cancers. Immune memory is a central feature of protective adaptive immunity, and mucosal barrier tissues are major sites of pathogen invasion. Yet, fundamental gaps remain in our knowledge of human T cell immune memory durability in mucosal barrier sites. Here, we employed minimally invasive nasal swab sampling to directly detect, and longitudinally assess, human antigen-specific T cells in two upper airway tissue sites (∼ 900 samples) and peripheral blood. Antigen-specific memory CD8 and CD4 T cells were detected in upper airway tissue of ≥ 95% of individuals. Both memory CD8 and CD4 T cells were sustained in both mucosal epithelial and mucosal lymphoid tissue over the course of 18+ months, with monthly sampling, and no clear evidence of decline. Virus-specific upper airway CD8 and CD4 T cells were predominantly resident memory T cells (T_RM_) throughout the 18+ month period of observation. These findings can inform future T cell vaccines and therapeutics.

---

The upper respiratory tract represents a key barrier site for the entry of respiratory pathogens. T cells in the upper airway may play important roles in controlling respiratory pathogens, but understanding of these T cells in humans, and whether they exhibit durable immune memory, has remained limited due to the technical challenges in mucosal sampling and detection of rare antigen-specific memory lymphocytes (*1–7*).

CD8 and CD4 T cells have been shown to play roles in mediating respiratory virus clearance and providing protection against respiratory virus infections (*8–13*). T cell cross-recognition and immunity may provide long-term immune protection even for respiratory viruses that escape humoral immunity via mutational evolution of neutralizing antibody escape variants (*1*, *10*, *14–21*). Tissue-resident memory T cells (T_RM_) are defined by long-term tissue retention and persistence in the absence of cognate antigen (*18*). T_RM_ share core expression profiles that enable persistence within tissues but display diverse functional phenotypes tailored to the specific tissue niches in which they reside (*2*, *3*, *19*). T_RM_ in mucosal barrier sites have been shown to provide enhanced protective immunity in local immune surveillance and defense in animal models (*7*, *18*, *19*, *22*). T cells can facilitate viral clearance by a range of mechanisms, including direct killing of infected cells, secretion of cytokines to drive antiviral resistance programs in tissues, recruitment of other cell types to the site of infection, and enhancement of local antibody concentrations, among other mechanisms of action (*1*, *23*). Virus-specific CD8 and CD4 T_RM_ phenotypes in the human upper airway remain incompletely defined (*8*, *17*, *24*). More importantly, the durability of virus-specific CD8 and CD4 memory T cells in human upper airway remains unknown, both for T_RM_ and any other T cell immune memory subsets that may be present, with key implications for the relevance of upper airway T cells in protective immunity.

Peptide-MHC (pMHC) multimers have been used in a few studies to track circulating antigen-specific T cell memory in human peripheral blood by direct binding of epitope-specific cells (*25*), including after primary SARS-CoV-2 (SARS2) infection or COVID-19 (COVID) vaccination (*26–29*), providing evidence of memory T cell longevity in blood, consistent with detection of circulating memory T cells in humans observed by other experimental approaches (*1*). CD4 T cells were assessed in human lymph node samples collected from individuals receiving an initial COVID mRNA vaccination series, without a history of SARS2 infection (*30*, *31*). SARS2-specific CD8 and CD4 T cells were detected in human upper airway tissue at a single timepoint in our prior report (*8*). Studies of human influenza T cell memory via direct detection with pMHC multimers have also primarily been restricted to T cells from peripheral blood, with limited assessment of longevity (*32*). Overall, direct detection (pMHC multimer binding) of antigen-specific mucosal T cells over time has not been performed for any human pathogen, for CD8 T cells or CD4 T cells, for mucosal epithelial tissue or mucosal lymphoid tissue. Thus, we aimed to address several of these critical knowledge gaps regarding human T cell memory.

## Respiratory virus-specific CD8 T cell memory is durable across blood and upper airway tissues

We enrolled a cohort of HLA-typed healthy adults (N = 34 total subjects. table S1-S2) to undergo longitudinal nasal swab and peripheral blood sampling (fig. S1A). Two distinct anatomic sites were targeted for nasal flocked swab sampling: the posterior nasopharynx (NP) and the mucosa of the inferior turbinate (MT), with the goal of collecting memory T cells from a secondary lymphoid organ (adenoid) and mucosal epithelium, respectively. We previously demonstrated that these two sites have distinct immune cell profiles, including differences in overall CD8 and CD4 T cell subset frequencies (*8*). Pertinent clinical data including respiratory virus infection and vaccination histories were collected at study entry and updated at each study visit. All participants reported a history of SARS2 infection and/or COVID vaccination prior to enrollment (table S1). A total of 457 study visits were conducted, collecting 1,010 total samples, over a period of 18+ months in 2024-2026.

Seven SARS2 class I peptide-MHC (pMHC-I) multimers (tetramers or pentamers) were selected to longitudinally track SARS2-specific memory CD8 T cells across time and tissues (Fig. 1A; fig. S1A, C-E; table S3). Peripheral blood mononuclear cell (PBMC) samples were collected approximately every 3 months (fig. S1A). Identification of circulating SARS2-specific CD8 memory T cells (T_mem_) by multiparametric spectral flow cytometry was performed by direct *ex vivo* multimer staining without enrichment or expansion (Fig. 1B-C; fig S1C,F-G). The specificity of pMHC-I multimer recognition by CD8 T_mem_ from HLA-matched participants was verified by dual staining (Fig. 1B-C, fig. S1F-G). Circulating CD8 T_mem_ responses to SARS2 spike (Fig. 1B) and non-spike (Fig. 1C; fig. S1F-G) epitopes were common among HLA-matched participants. Circulating SARS2-specific CD8 T_mem_ were durable over the ∼14-month blood sampling period (Fig. 1B-C; fig. S1F-G). All participants with relevant class I MHC alleles had detectable circulating CD8 T cell memory to one or more SARS2 pMHC-I multimers (Fig. 1H). The majority of participants showed no serologic evidence of SARS2 re-exposure based on plasma SARS2-nucleocapsid (N) and spike (S)-specific IgG (fig. S1B), indicating that detection of circulating SARS2-specific CD8 T cell memory was not attributable to frequent re-infections.

**Figure 1.**
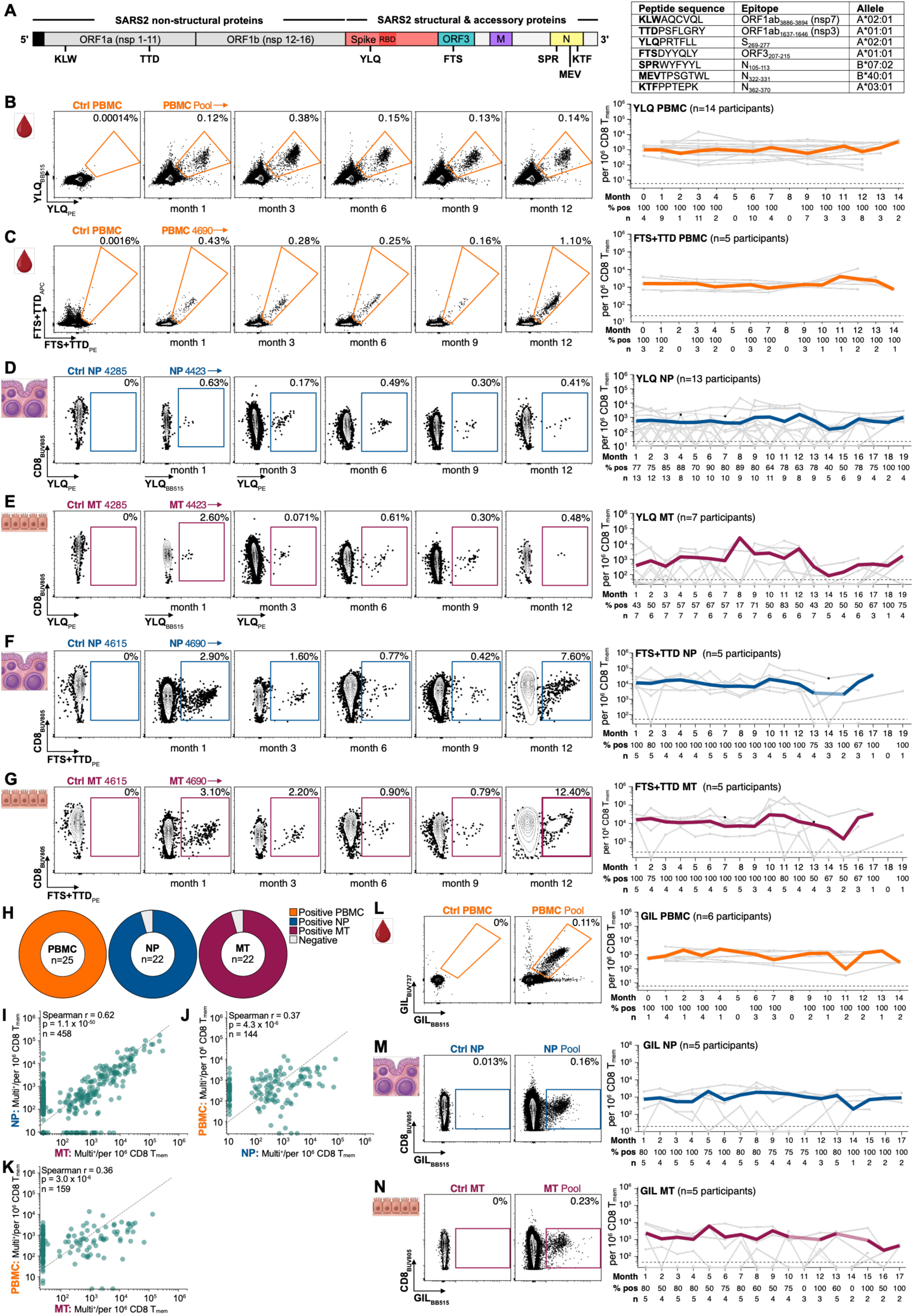
CD8 T cell memory is durable in blood and upper airway tissues. A. Left: Simplified SARS2 genome annotated with the class I pMHC multimers utilized in this study to identify SARS2-specific CD8 memory T cells (shorthand multimer naming = first three amino acids of the peptide sequence). Receptor binding domain (RBD) of spike in darker red. Right: Corresponding full length amino acid peptide sequences, viral epitopes, and HLA restrictions. See Table S3 for additional details. B. Left: Concatenated longitudinal flow cytometry data for PBMC from HLA-matched subjects dual stained with the YLQ HLA-A*02:01-restricted SARS2 spike multimer (n = 3-4 subjects; one subject missed the 3-month blood draw). Negative control (Ctrl) = unstained memory CD8 T cells from the same subjects. Right: Longitudinal circulating YLQ-specific memory CD8 T cell responses for n = 14 HLA-matched subjects. Thin gray lines = responses for individual subjects; solid gray dots = positive responses. Thick colored line = group geomean. Dashed black line = threshold for positivity. % pos = percent positive responders for each time point. C. Left: Example longitudinal flow cytometry plot examples of dual staining of PBMC from a single HLA-matched subject (4690) with the paired HLA-A*01:01-restricted non-spike FTS and TTD pentamers. Negative control (Ctrl) = unstained memory CD8 T cells from the same subject. Right: Longitudinal circulating FTS+TTD-specific memory CD8 T cell responses for n = 5 HLA-matched subjects. D. Left: Example flow cytometry plots of YLQ multimer-stained NP swab from an HLA-mismatched subject (Ctrl; 4285) and longitudinal YLQ-stained NP swabs from an HLA-A*02:01 heterozygous subject (4423). Right: Longitudinal YLQ responses in NP swabs from HLA-A*02:01 positive subjects (n = 13). Only subjects with at least 3 positive YLQ responses (> or = 2 multimer-positive cells and > 1,000 CD8 T cells/swab) were included in the longitudinal NP group plot. Thin gray lines = responses for individual subjects (dashed gray lines connect data points when visits were skipped or swabs failed QC); solid gray dots = positive responses from swabs with > 1,000 CD8 T cells, solid black dots = positive responses from swabs with < 1,000 CD8 T cells, open gray circles = negative responses from swabs with > 1,000 CD8 T cells. Thick colored line = group geomean; only solid gray dots contributed to geomean calculations. Dashed black line = threshold for positivity. E. Left: Example flow cytometry plots of a YLQ multimer-stained MT swab from the same HLA-mismatched subject (Ctrl) and longitudinal YLQ-stained MT swabs from the same HLA-A*02:01 positive subject as in D. Right: Longitudinal YLQ responses in MT swabs from HLA-A*02:01 positive subjects as D. Only subjects with at least 3 positive YLQ responses (> or = 2 multimer-positive cells and > 100 CD8 T cells/swab) were included in the longitudinal MT group plot (n = 7). Thin gray lines = responses for individual subjects (dashed gray lines connect data points when visits were skipped or swabs failed QC); solid gray dots = positive responses from swabs with > 100 CD8 T cells, open gray circles (floored) = negative responses from swabs with > 100 CD8 T cells. Thick colored line = group geomean; only solid gray dots contributed to geomean calculations. Dashed black line = threshold for positivity. F. Left: Example flow cytometry plots of FTS+TTD pentamer-stained CD8 memory T cells from the NP swab of an HLA-mismatched subject (Ctrl; 4615) and from longitudinal NP swabs from an HLA-A*01:01 positive subject (4690). Right: Longitudinal FTS+TTD responses in NP swabs from HLA-A*01:01 positive subjects. Only subjects with at least 3 positive FTS+TTD responses (> or = 2 pentamer-positive cells and > 1,000 CD8 T cells/swab) were included in the longitudinal NP group plot (n = 5). G. Left: Example flow cytometry plots of an FTS+TTD pentamer-stained MT swab from the same HLA-mismatched subject (ctrl; 4615) as in F, and longitudinal FTS+TTD-stained MT swabs from the same HLA-A*01:01 positive subject (4690) as in F. Right: Longitudinal FTS+TTD responses in MT swabs from the same HLA-A*01:01 positive subjects as F. Only subjects with at least 3 positive FTS+TTD responses (> or = 2 pentamer-positive cells and > 100 CD8 T cells/swab) were included in the longitudinal MT group plot (n = 5). H. Proportion of study subjects with relevant HLA alleles with responses to at least 1 SARS2 class I multimer in PBMC, NP, and MT swabs. n = unique subjects who provided each sample type for multimer staining; 3 subjects only provided blood samples for PBMC testing. Positive = samples that met QC metrics and had positive multimer responses. Negative = samples that met QC metrics but had negative responses. See Methods for details. I. Correlations between SARS2-specific memory CD8 T cell frequencies for all SARS2 class I multimers for matched NP and MT swab samples (n = 458) from the same subjects and time points stained with the same multimer(s). Zero values were set to the minimum threshold for positivity for all SARS2 class I multimers by site (NP, MT). J. Correlations between SARS2-specific memory CD8 T cell frequencies for all SARS2 class I multimers for matched NP swab and PBMC samples (n = 144) from the same subjects and time points stained with the same multimer(s). Zero values were set to the minimum threshold for positivity for all SARS2 class I multimers by site (NP, PBMC). K. Correlations between SARS2-specific memory CD8 T cell frequencies for all SARS2 class I multimers for matched MT swab and PBMC samples (n = 159) from the same subjects and time points stained with the same multimer(s). Zero values were set to the minimum threshold for positivity for all SARS2 class I multimers by site (MT, PBMC). L. Left: Example flow cytometry plots of concatenated memory CD8 T cell responses in dual Influenza A class I multimer (GIL)-stained PBMC from HLA-A*02:01 positive subjects (n = 6). Negative control (Ctrl) = concatenated data for unstained memory CD8 T cells from the same subjects. Right: Longitudinal circulating GIL-specific memory CD8 T cell responses for n = 6 HLA-matched subjects. M. Left: Example flow cytometry plots of concatenated memory CD8 T cell responses in GIL-stained NP swabs from pooled HLA-mismatched (Ctrl) and HLA-A*02:01 positive subjects. Right: Longitudinal GIL responses in NP swabs from HLA-A*02:01 positive subjects. Only subjects with at least 3 positive GIL responses (> or = 2 pentamer-positive cells and > 1,000 CD8 T cells/swab) were included in the longitudinal NP group plot (n = 5). N. Left: Example flow cytometry plots of concatenated memory CD8 T cell responses in GIL-stained MT swabs from the same HLA-mismatched (Ctrl) and HLA-A*02:01 positive subjects as in M. Right: Longitudinal GIL responses in MT swabs from the same HLA-A*02:01 positive subjects as M. Only subjects with at least 3 positive GIL responses (> or = 2 pentamer-positive cells and > 100 CD8 T cells/swab) were included in the longitudinal MT group plot (n = 5).

We examined the durability of upper airway CD8 T cell memory in the same cohort using the same pMHC-I multimers. Longitudinal NP and MT swab sampling was performed monthly. Upper airway SARS2-specific CD8 T_mem_ were detectable by direct multimer staining of MT and NP swab samples without enrichment or expansion (Fig. 1D-K, fig. S1F-G). Remarkably, upper airway SARS2 spike-specific CD8 T_mem_ were persistently detectable in the majority of monthly NP and MT swab samples collected over the course of >18 months (Fig. 1D-E). Similar results were also observed for SARS2-specific CD8 T_mem_ cells recognizing non-spike antigens in upper airway swab samples (Fig. 1F-G; fig. S1F-G). Overall, 95% of participants with relevant class I MHC alleles had detectable NP and MT CD8 T cell memory (Fig. 1H), and those SARS2-specific CD8 T_mem_ could be detected across time (Fig. 1D-G, fig. S1F-G). No clear evidence of decline in CD8 T_mem_ cell frequencies was observed over ∼18 months in NP or MT tissues (Fig. 1D-G, fig. S1F-G).

Relationships between the frequencies of antigen-specific CD8 T_mem_ across anatomic sites were examined. Virus-specific CD8 T_mem_ strongly correlated between upper airway mucosa (MT) and upper airway lymphoid tissue (NP) (Fig. 1I; Spearman r = 0.62 and p = 1 x 10^-50^). In contrast, circulating and upper airway CD8 T_mem_ frequencies correlated more weakly (Fig. 1J and 1K; Spearman r = 0.37 and 0.36 for PBMC and NP or MT, respectively. p = 4 x 10^-6^ and 3 x 10^-6^).

The study was expanded to examine CD8 T cell memory to a non-SARS2 pathogen. Influenza A pMHC-I multimer-specific CD8 T_mem_ cells were detectable for >12 months in PBMC (Fig. 1L), NP swab (Fig. 1M), and MT swab samples (Fig. 1N). Thus, durable upper airway CD8 T cell memory in humans is not exclusive to SARS2.

## Respiratory virus-specific CD4 T cell memory is durable across blood and upper airway tissues

Six SARS2 class II peptide-MHC (pMHC-II) multimers were selected to directly identify SARS2-specific CD4 T_mem_ across time and tissues (Fig. 2A, fig. S2A-C, table S3) in the same HLA-typed study cohort (fig. S1A, tables S1-S2). The majority of participants were HLA-DPB1*04:01 or *04:02 positive (table S2). Dual multimer staining of PBMC was performed to verify the specificity of pMHC-II multimer recognition by CD4 T_mem_ from a series of HLA-matched participants (Fig. 2B-C; fig. S2D-F). Circulating spike (Fig. 2B; fig. S2D-E) and non-spike (Fig. 2C, fig. S2F) SARS2-specific CD4 T_mem_ cells were commonly detected. SARS2-specific memory CD4 T cell frequencies were durable over the ∼14-month blood sampling period, with no evidence of a half-life (Fig. 2B-C; fig. S2G-I).

**Figure 2.**
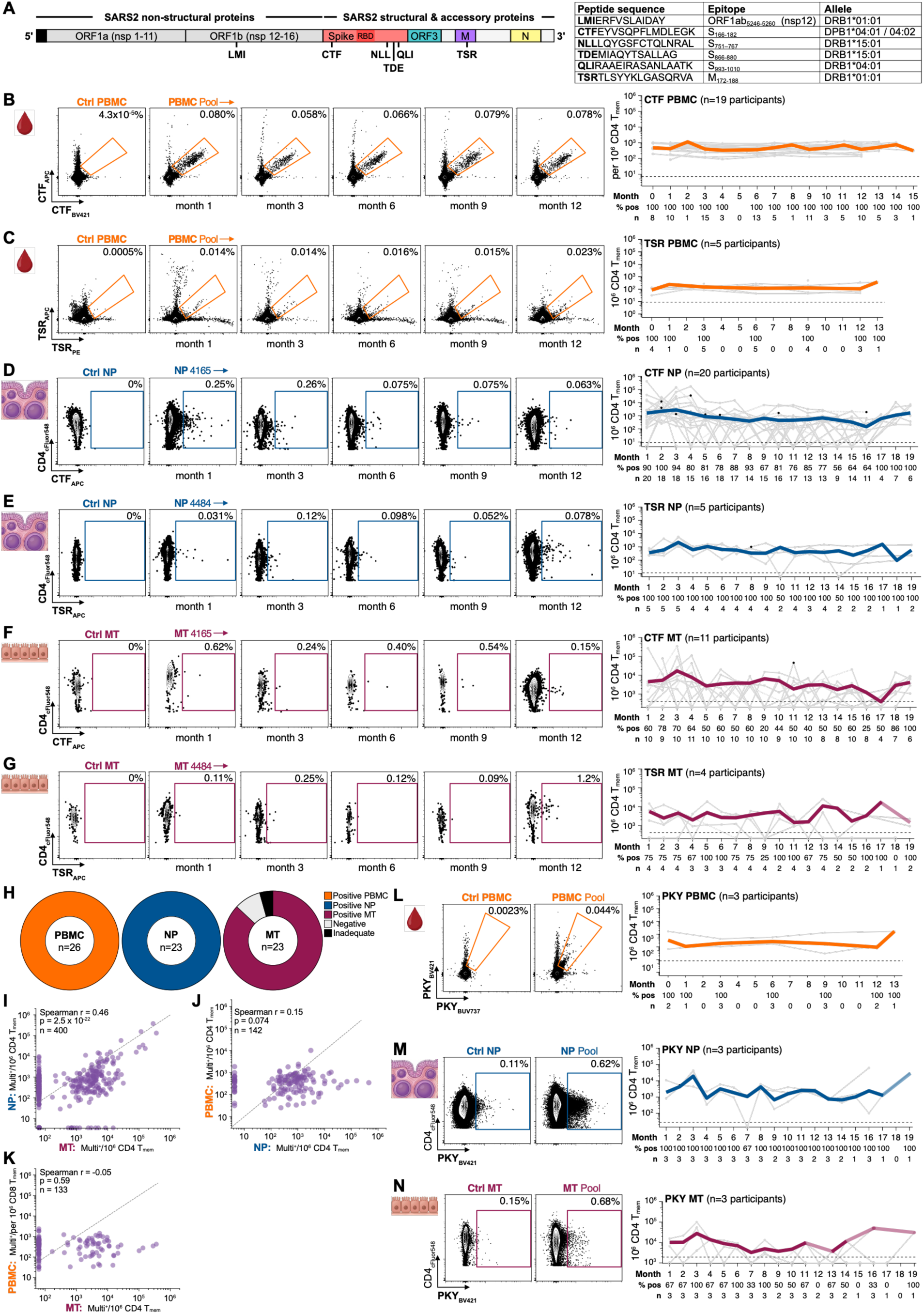
CD4 T cell memory is durable across blood and upper airway tissues. A. Left: Simplified SARS2 genome annotated with the class II pMHC multimers utilized in this study to identify SARS2-specific CD4 memory T cells (shorthand multimer naming = first three amino acids of the peptide sequence). RBD of spike in darker red. Right: Corresponding full length amino acid peptide sequences, viral epitopes, and HLA restrictions. See Table S3 for additional details. B. Left: Concatenated longitudinal flow cytometry data for PBMC from HLA-matched subjects that were dual stained with the CTF DPB1*04:01/04:02-restricted SARS2 spike multimer (n = 4 subjects). Negative control (Ctrl) = unstained memory CD4 T cells from the same subjects. Right: Longitudinal circulating CTF-specific memory CD4 T cell responses for n = 19 HLA-matched subjects. C. Left: Concatenated longitudinal flow cytometry data for PBMC from HLA-matched subjects that were dual stained with the TSR DRB1*01:01-restricted non-spike multimer (n = 4 subjects). Negative control (Ctrl) = unstained memory CD4 T cells from the same subjects. Right: Longitudinal circulating TSR-specific memory CD4 T cell responses for n = 5 HLA-matched subjects. D. Left: Example flow cytometry plots of a CTF multimer-stained NP swab from an HLA-mismatched subject (Ctrl) and from longitudinal CTF-stained NP swabs from a DPB1*04:01/*04:02-positive subject (4165). Right: Longitudinal CTF responses in NP swabs from DPB1*04:01/*04:02-positive subjects. Only subjects with at least 3 positive CTF responses (> or = 2 multimer-positive cells and > 1,000 CD4 T cells/swab) were included in the longitudinal NP group plot (n = 20). Thin gray lines = responses for individual subjects (dashed gray lines connect data points when visits were skipped or swabs failed QC); solid gray dots = positive responses from swabs with > 1,000 CD4 T cells, solid black dots = positive responses from swabs with < 1,000 CD4 T cells, open gray circles = negative responses from swabs with > 1,000 CD4 T cells. Thick colored line = group geomean; only solid gray dots contributed to geomean calculations. Dashed black line = threshold for positivity. E. Left: Example flow cytometry plots of a TSR multimer-stained NP swab from an HLA-mismatched subject (Ctrl) and from longitudinal TSR-stained NP swabs from an DRB1*01:01 positive subject (4484). Right: Longitudinal TSR responses in NP swabs from DRB1*01:01 positive subjects (n = 5). Only subjects with at least 3 positive TSR responses (> or = 2 multimer-positive cells and > 1,000 CD4 T cells/swab) were included in the longitudinal NP group plot. F. Left: Example flow cytometry plots of a CTF multimer-stained MT swab from an HLA-mismatched subject (Ctrl) and from longitudinal CTF-stained MT swabs from the same DPB1*04:01/*04:02-positive subject (4165) as in D. Right: Longitudinal CTF responses in MT swabs from DPB1*04:01/*04:02-positive subjects. Only subjects with at least 3 positive CTF responses (> or = 2 multimer-positive cells and > 1,000 CD4 T cells/swab) were included in the longitudinal MT group plot (n = 11). Thin gray lines = responses for individual subjects (dashed gray lines connect data points when visits were skipped or swabs failed QC); solid gray dots = positive responses from swabs with > 100 CD4 T cells, solid black dots = positive responses from swabs with < 100 CD4 T cells, open gray circles = negative responses from swabs with > 100 CD4 T cells. Thick colored line = group geomean; only solid gray dots contributed to geomean calculations. Dashed black line = threshold for positivity. G. Left: Example flow cytometry plots of a TSR multimer-stained MT swab from an HLA-mismatched subject (Ctrl) and longitudinal CTF-stained MT swabs from the same DRB*01:01 positive subject (4484) as in D. Right: Longitudinal TSR responses in MT swabs from DRB1*01:01 positive subjects. Only subjects with at least 3 positive TSR responses (> or = 2 multimer-positive cells and > 100 CD4 T cells/swab) were included in the longitudinal MT group plot (n = 4). H. Proportion of study subjects with relevant HLA alleles with responses to at least 1 SARS2 class II multimer in PBMC, NP, and MT swabs. n = unique subjects who provided each sample type for multimer staining; 3 subjects only provided blood samples for PBMC testing. Positive = samples that met QC metrics and had positive multimer responses. Negative = samples that met QC metrics but had negative responses. Inadequate = samples that failed to meet QC metrics. See Methods for details. I. Correlations between SARS2-specific memory CD4 T cell frequencies for all SARS2 class II multimers for matched NP and MT swab samples (n = 400) from the same subjects and time points stained with the same multimer(s). Zero values were set to the minimum threshold for positivity for all SARS2 class II multimers by site (NP, MT). J. Correlations between SARS2-specific memory CD4 T cell frequencies for all SARS2 class II multimers for matched NP swab and PBMC samples (n = 142) from the same subjects and time points stained with the same multimer(s). Zero values were set to the minimum threshold for positivity for all SARS2 class II multimers by site (NP, PBMC). K. Correlations between SARS2-specific memory CD4 T cell frequencies for all SARS2 class II multimers for matched MT swab and PBMC samples (n = 133) from the same subjects and time points stained with the same multimer(s). Zero values were set to the minimum threshold for positivity for all SARS2 class II multimers by site (MT, PBMC). L. Left: Example flow cytometry plots of concatenated memory CD4 T cell responses in dual Influenza A class II multimer (PKY)-stained PBMC from DRB1*07:01 positive subjects (n = 3). Negative control (Ctrl) = concatenated unstained memory CD4 T cell data from the same subjects. Right: Longitudinal circulating PKY-specific memory CD4 T cell responses for n = 3 HLA-matched subjects. M. Left: Example flow cytometry plots of concatenated memory CD4 T cell responses in PKY-stained NP swabs from HLA-mismatched (Ctrl) and DRB1*07:01 positive subjects. Right: Longitudinal PKY responses in NP swabs from DRB1*07:01 positive subjects. Only subjects with at least 3 positive PKY responses (> or = 2 pentamer-positive cells and > 1,000 CD4 T cells/swab) were included in the longitudinal NP group plot (n = 3). N. Left: Example flow cytometry plots of concatenated memory CD4 T cell responses in PKY-stained MT swabs from the same HLA-mismatched control (Ctrl) and DRB1*07:01positive subjects as in M. Right: Longitudinal PKY responses in MT swabs from the same DRB1*07:01 positive subjects as M. Only subjects with at least 3 positive PKY responses (> or = 2 pentamer-positive cells and > 100 CD4 T cells/swab) were included in the longitudinal NP group plot (n = 3).

Upper airway SARS2-specific CD4 T cell memory was assessed in NP and MT tissue over time by monthly longitudinal swab sampling. Notably, SARS2 spike-specific CD4 T_mem_ were present in NP tissue and sustained for >18 months, with no evidence of a half-life (Fig. 2D; fig. S2J-K). Non-spike-specific CD4 T cell memory was also observed in NP tissue (Fig. 2E, fig. S2L). Durable SARS2 upper airway CD4 T cell memory was observed for both adenoid and non-adenoid NP swab samples (Fig. S2M; Methods). Fewer CD4 T cells are present in MT swabs compared to CD8 T cells (CD8:CD4 ratio ∼5:1) (*8*). Nevertheless, SARS2-specific memory CD4 T cells were directly identifiable in MT swabs by spectral flow cytometry without enrichment or expansion (Fig. 2F-G, fig. S2N-P). SARS2-specific CD4 T_mem_ were detectable in MT mucosa for >18 months (Fig. 2F-G, Fig. S2N-O).

Overall, 100% of participants with the relevant MHC class II alleles had detectable circulating SARS2-specific CD4 T_mem_, 95% of participants had detectable SARS2-specific CD4 T_mem_ in NP swabs, and 91% of participants had detectable SARS2-specific CD4 T_mem_ in MT swabs (Fig. 2H). In each case, no obvious decline in CD4 T cell memory was detected over time (Fig. 2B-G; fig. S2G-L, N-O). Relationships between CD4 T_mem_ across anatomic sites were examined (Fig. 2I-K). Frequencies of SARS2-specific CD4 T_mem_ were positively correlated between MT and NP tissue sites (Fig. 2I; Spearman r = 0.46, p = 2.5 x 10^-22^). Notably, no significant correlation was found between antigen-specific CD4 T_mem_ in blood and upper airway NP tissue (Fig. 2J) or MT mucosa (Fig. 2K).

Upper airway CD4 T cell memory to a different pathogen was assessed using an HLA-DRB1*07:01 restricted Influenza A specific pMHC-II multimer (PKY) (table S3). Influenza A-specific CD4 T_mem_ were detected for > 12 months in PBMC (Fig. 2L) and > 18 months in NP (Fig. 2M) and MT (Fig. 2N) swab samples from HLA-matched subjects (table S2), demonstrating that long-lasting pathogen-specific upper airway CD4 T cell memory is not exclusive to SARS2.

## Diversity of bulk and antigen-specific upper airway CD8 T cell subsets

To better characterize MT and NP CD8 T_RM_, we developed and optimized an expanded T cell-centric spectral flow cytometry immunophenotyping panel to evaluate antigen-specific and bulk upper airway CD8 T_RM_ and other CD8 T cell subsets. Nearly 400 MT and 400 NP monthly swabs in total, from 31 participants, were examined by detailed immunophenotyping. Of these, 23 participants provided longitudinal MT and NP swabs for > 12 months. Upper airway memory CD8 T cell phenotypes in MT and NP tissue sites were distinct from memory CD8 T cells in blood (Fig. 3A-C, fig. S3A). Over 90% of CD8 T_mem_ in both MT and NP tissues expressed CD69 (Fig. 3A). The great majority of the CD69^+^ CD8 T cells in both tissues co-expressed the canonical T_RM_ marker, CD103 (Fig. 3A). CXCR6 was also co-expressed by the majority of MT and NP CD8 T_mem_ (Fig. 3A-C). Frequencies of CXCR6^+^ CD8 T cells were stable over time in both tissue sites (Fig. 3B). Other potential T_RM_ markers such as CD49a (Fig. 3A) were less frequently expressed by MT and NP CD8 T_mem_, but still stable over time (fig. S3A). Overall, upper airway CD8 T_mem_ in MT and NP tissue shared similar immunophenotypes, despite deriving from nasal mucosal epithelium versus adenoid lymphoid tissue sampling sites.

**Figure 3.**
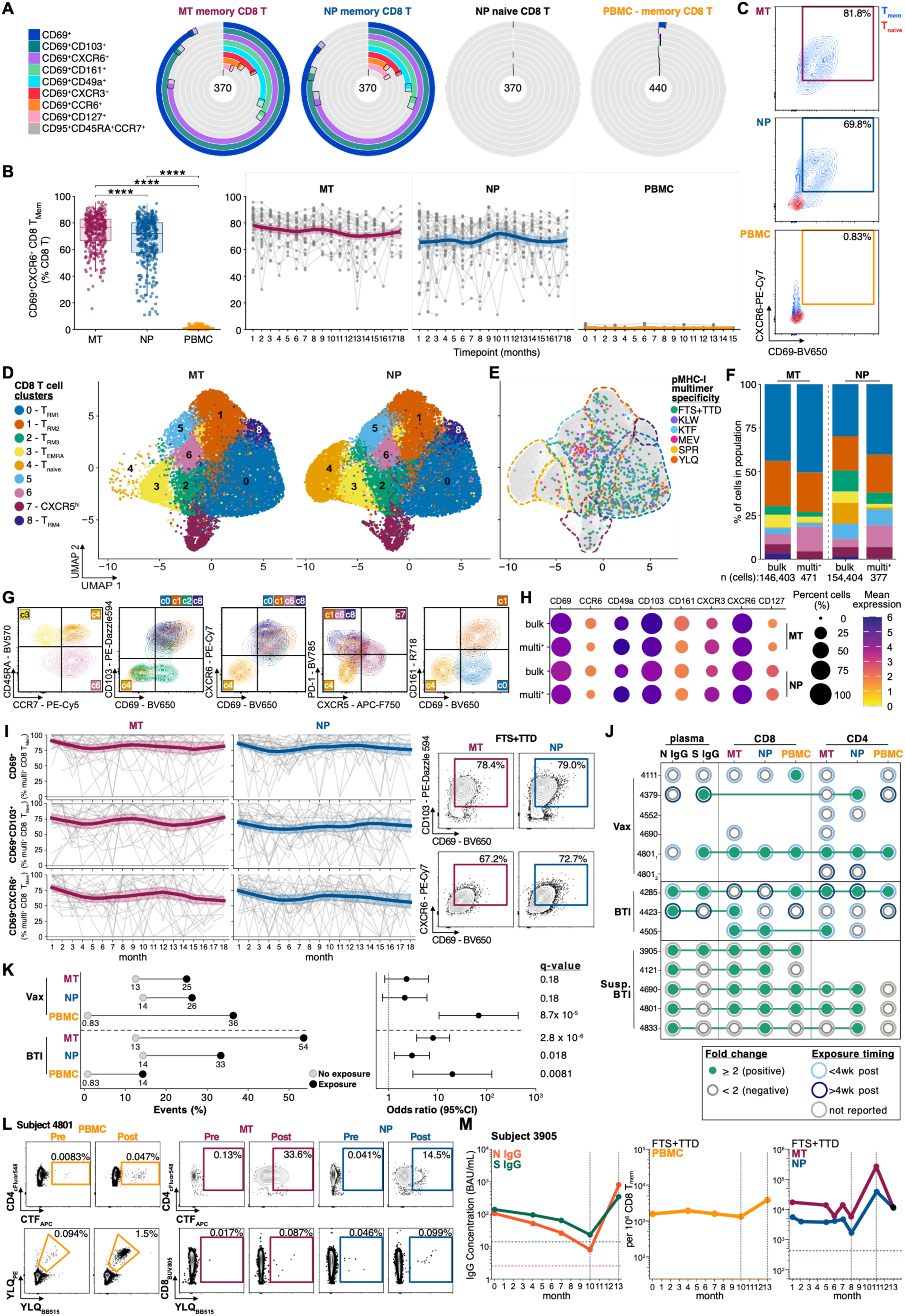
Upper airway CD8 T cells share conserved features and antigen-specific T cells respond to local antigen exposure. A. ‘Belt’ plots with concentric rings demonstrating the proportion of memory CD8 T cells from MT swabs, NP swabs, or PBMC and naive CD8 T cells from NP swabs that express CD69 alone or in combination with other putative T_RM_ markers (values in the centers of the concentric rings = number of samples); medians and 95% confidence intervals (CI) are shown. B. Cross-sectional (left) and longitudinal (right) co-expression of CD69 and CXCR6 by non-naive CD8 T cells in MT and NP swabs, PBMC. Box and whisker plots display median, interquartile range (IQR), and range. Left: Pairwise Wilcoxon tests with Bonferroni correction for multiple comparison testing; **** = p-value < or = 0.0001. C. Example flow cytometry contour plots of CD69 and CXCR6 co-expression by non-naive CD8 T cells in MT swabs (blue, T_Mem_; CD45RA^-^ CCR7^-^, CD45RA^-^CCR7^+^, and CD45RA^+^CCR7^-^). Overlaid naive (red, CD45RA^+^CCR7^+^, T_naive_) versus non-naive (blue, T_Mem_) CD8 T cells in NP swabs and PBMC. % CD69^+^CXCR6^+^ of parental (non-naive CD8 T cells) is shown. D. Seurat UMAPs split by anatomic site (MT and NP) of single cell CD8 T cell flow cytometry data from 740 MT and NP samples from 31 subjects. Random downsampling prior to Seurat object construction with up to 5,000 cells per subject and site yielded a Seurat object with 307,121 CD8 T cells. Fourteen surface markers were used as variable features: CD69, CD103, CD49a, CXCR6, CCR6, CD161, CXCR3, CD127, CXCR5, CCR7, CD45RA, PD-1, CD25, and CD95. The result was a total of 9 CD8 T cell clusters at a resolution of 0.2. For visualization, 50,000 cells were randomly subsampled from each object. See Methods and Fig. S3C for additional details. E. Seurat UMAP of the combined downsampled MT and NP data from D as a background of gray, multimer-negative CD8 T cells with an overlay of multimer-positive CD8 T cells from 23 HLA-matched subjects color-coded by class I multimer(s)-specificity. Cells binding more than one multimer (other than those binding FTS/TTD, where the allele restriction is shared and binding to FTS vs TTD could not be deconvoluted) were excluded. To aid visualization, approximate outlines of the clusters from D are shown in the corresponding colors and SARS2 multimer-positive cells are plotted at an increased point size relative to the background cells; n = 848 pMHC-multimer-positive CD8 T cells are shown, of ∼15,000 total pMHC-multimer-positive CD8 T cell events collected. F. Stacked bar plot showing (from left to right) the proportion of bulk MT, multimer-positive MT, bulk NP, and pMHC-multimer-positive NP CD8 T cells contributing to each of the 9 UMAP clusters shown in D; n = the total number of bulk and antigen-specific CD8 T cells included in the analysis. G. Flow cytometry plots generated from the CD8 T cells assigned to each Seurat cluster demonstrating differential marker expression by the CD8 T cell clusters related to naive (CD45RA^+^CCR7^+^) vs memory, T_RM_ (CD69, CD103, CXCR6), CXCR5^hi^, and other CD8 T cell subset phenotypes. H. Dot plot with unscaled surface protein expression of CD69 and other putative T_RM_ markers by bulk MT and NP and SARS2 class I multimer-positive MT and NP CD8 T cells. Dot size = % of total CD8 T cells expressing CD69 or % of total CD69^+^ CD8 T cells expressing each additional marker per sample; averaged across samples per site for bulk cells (MT ∼1.5 and NP ∼3.7 million CD8 T cells) from flow cytometry data, or averaged first across epitope specificities per subject per timepoint then across subjects per site for multimer-positive cells (MT 4,907 and NP 10,675 total pMHC-I multimer-positive CD8 T cells), with a minimum of 3 CD69^+^ cells per sample required. Dot color = mean asinh fluorescence intensity (cofactor 150, fixed color scale 0–6, dark = high expression) computed from the projected multimer-positive cell Seurat object subset (15,279 CD8 T cells) or from the Seurat object for bulk CD8 T cells. I. Left: T_RM_ marker (CD69, CD103, CXCR6) expression stability in SARS2 class I multimer-positive memory CD8 T cells over time by flow cytometry. % of pooled multimer-positive CD8 memory T cells in MT (n = 232) or NP (n = 280) swabs expressing each indicated marker. Data from swabs collected monthly for up to 18 months from up to 30 subjects per plot (individual timepoints include 6–25 subjects). Thin gray lines represent individual subject’s trajectories. Thick colored line = LOWESS-smoothed mean; shaded band indicates ±SEM. Right: Example concatenated flow cytometry data of CD69 and CD103 co-expression (top) and CD69 and CXCR6 co-expression for FTS+TTD pMHC-I multimer positive memory CD8 T cells in n = 71 MT and 71 NP swabs. J. Summary of reported COVID vaccinations (Vax) and SARS2 breakthrough infections (BTI), and suspected infections (Susp. BTI) that occurred during the study. Associated fold-change post-event in SARS2-specific plasma IgG (S = WT spike, N = nucleocapsid) and multimer responses in MT, NP, PBMC samples are indicated. For Vax, only spike-specific pMHC responses were evaluated. For reported events, outer rings indicate whether samples were collected < 4 weeks (light blue) or > 4 weeks (dark blue) post-event; based on reported vaccination or BTI symptom onset dates. For suspected infections (Susp. BTI), outer rings are light gray indicating no reported symptom onset date. Inner circles indicate whether the two-fold increase threshold was met: positive = filled green, negative = unfilled dark gray. Blanks indicate that data was available for pre-/post-event fold change analyses. K. Left: The multimer responses associated with reported vaccination (Vax) or reported/suspected SARS2 infection (BTI) events from I were examined, and the proportion of reported/suspected Ag-exposure events (“exposure” black dots on FC plot) that resulted in a four-fold or greater increase in SARS2 multimer-specific memory T cell frequencies in MT, NP, and PBMC samples was calculated. The proportion of consecutive sampling time points with four-fold or greater increases in SARS2 multimer-specific memory T cell frequencies for time points without any reported/suspected events was also calculated (“no exposure” gray dots). Right: Fisher’s exact testing was performed between the exposure and no exposure groups, and the resulting odds ratios with 95% confidence intervales and q-values following Benjamini-Hochberg correction for multiple comparisons are shown. See Methods, Table S1, and Fig. S3F-G for additional details. L. Example flow cytometry data from pre- (month 1) and post-COVID mRNA vaccination (month 3) for spike-specific pMHC-I (YLQ) and pMHC-II (CTF) multimer responses pre- versus post-vaccination in PBMC, MT, and NP samples for the first booster vaccination reported by subject 4801(4801_1_ in Fig. 3J). L. Longitudinal SARS2 serology and multimer response data from subject 3905 (Susp. BTI in Fig. 3J), illustrating a suspected SARS2 BTI that resulted in concomitant increases in SARS2 IgG (S = WT spike, N = nucleocapsid) and pMHC-I FTS+TTD multimer responses in PBMC, MT and NP swabs. Horizontal dashed lines indicate the limit of detection for IgG (green for S and orange for N) or threshold for positivity for multimers (red = MT, blue = NP, light orange = PBMC). Vertical dotted lines indicate the time period corresponding to the suspected exposure based on the rise in N IgG. Serology was performed using plasma from the same peripheral blood samples as PBMC. This subject provided a blood sample prior to the start of MT and NP swab sampling; blood sampling months 10-13 correspond to swab sampling months 8-11.

MT and NP CD8 T_RM_ phenotypes were further explored via unsupervised analyses of upper airway swab high-dimensional spectral flow cytometry data. Several CD8 T_mem_ cell phenotypic clusters were observed across MT and NP sites, including multiple CD8 T_RM_ clusters (Fig. 3D, fig. S3B-C). High expression of CD161 was a distinguishing feature of T_RM_ cluster 1 (T_RM1_), and high CXCR5 expression was a prominent feature of cluster 7 (Fig. 3G, fig. S3B-C). SARS2-specific CD8 T_mem_ were distributed across the majority of the Seurat clusters (Fig. 3E-F). Subtle differences were observed in SARS2-specific versus bulk upper airway CD8 T cell distributions across Seurat clusters (Fig. 3F) and by individual marker expression (Fig. 3H). SARS2-specific CD8 T_RM_ were enriched for CXCR3 expression, consistent with their genesis in an antiviral response (Fig. 3H). The majority of SARS2-specific CD8 T_mem_ in MT and NP tissue sites were T_RM_ based on expression of CD69, CD103, and CXCR6 (Fig. 3E-H). Each of these T_RM_ markers appeared to be maintained consistently over time for SARS2-specific CD8 T_mem_ (Fig. 3I). Class I influenza A multimer-specific MT and NP CD8 T_mem_ were similarly observed to predominantly be phenotypically CD8 T_RM_ (fig. S3D). Overall, these findings indicate that CD8 T cell persistence in both upper airway non-lymphoid mucosal and lymphoid tissue sites is predominantly associated with a T_RM_ program.

## Antigen-specific memory T cell responses following vaccination or infection

Six individuals reported receiving one or more COVID mRNA vaccine booster immunization(s) during the study (table S1). Spike pMHC-II-specific T cells have been observed in peripheral blood and draining lymph nodes for up to 6 months post-vaccination (*30*, *31*). We assessed whether differences in upper airway SARS2 spike-specific T cell memory could be observed pre- and post-vaccination. Paired pre-/post-vaccination SARS2 spike pMHC multimer-stained MT and NP swab data was available for six vaccination events, with temporally paired peripheral blood data available in some cases (Fig. 3J). Vaccination resulted in a ≥ 2-fold increase in circulating spike IgG in 2 of 3 cases, and vaccination resulted in a ≥ 2-fold increase in circulating SARS2 spike-specific T cells in 2 of the 3 events for which PBMC samples were available (Fig. 3J-K; fig. S3E-F). Cross-tissue increases in circulating and upper airway CD8 and CD4 T_mem_ were observed in only one case (4801_1_), and that subject coincidentally was sampled at an acute timepoint, 11 days post-vaccination (Fig. 3L).

We next evaluated whether local antigen exposure from SARS2 infection altered upper airway SARS2 T cell immunity. During the study, three participants reported symptomatic SARS2 breakthrough infections (BTI) with positive viral confirmatory testing (table S1). Pre-/post-BTI peripheral blood and nasal swab samples were available for two events (4285, 4423), and nasal swab samples were available for the third (4505). Increased plasma N IgG confirmed the two BTIs with plasma samples available (Fig. 3J, fig. S3E). It was then assessed whether confirmed BTI events were associated with increases in upper airway T cells. All three subjects with confirmed BTI had increased SARS2 pMHC-specific T_mem_ in MT swabs post-infection (Fig. 3J, fig. S3G). Increased SARS2-specific T_mem_ were also detected in NP swabs post-infection in 2 of the 3 confirmed BTIs (Fig. 3J, fig. S3G). Notably, only one of the two confirmed BTIs with PBMC samples resulted in increased circulating SARS2-specific T_mem_ (Fig. 3J, fig. S3G).

Using the confirmed BTI immunological outcomes as a benchmark, it was then assessed whether additional unreported/undetected SARS2 infections had occurred in this cohort. Five additional suspected SARS2 infections were identified based on an increase in plasma N IgG (Fig. 3J, fig. S3E). Notably, in each of the 5 cases of suspected infection, a concurrent increase in SARS2-specific T_mem_ in NP swabs was observed (Fig. 3J, fig. S3G). Furthermore, an increase in SARS2-specific T_mem_ in the MT site was also observed for 80% (4 of 5) of suspected infections (Fig. 3J, fig. S3G). While increases in SARS2-specific upper airway CD8 T_mem_ responses were observed in 100% of suspected infection cases, increases in circulating CD8 T_mem_ were only observed in 60% (3 of 5) (Fig. 3J, M; fig. S3G). For CD4 T cells, pMHC-II reagents were available for three subjects; mucosal SARS2-specific CD4 T_mem_ increases were observed in 100% (3 of 3) of those subjects, while none of those subjects (0 of 3) had any detectable CD4 T_mem_ changes in blood (Fig. 3J; fig. S3G).

Comparing changes in antigen-specific T cell frequencies across time and tissues, reported and suspected infection events were associated with significant increases (≥ 4-fold) in upper airway SARS2-specific T_mem_ (MT, *q* = 2.8 x 10^-6^. NP, *q* = 0.018. Fig. 3K). The full cohort represented ∼450 person-months of study time. There were eight reported/suspected SARS2 infection events, representing one event per ∼56 person-months. The majority of participants in this cohort exhibited no measurable infection event during the ∼18-month course of the study based on SARS2-specific plasma IgG serology or SARS2-specific circulating and upper airway memory CD8 and CD4 T cells. Based on the data collected, it appears that for most individuals SARS2-specific CD8 and CD4 T cell memory in blood, upper airway lymphoid tissue (NP), and upper airway non-lymphoid (MT) tissue is durable in the absence of measurable infections.

## Diversity of bulk and antigen-specific upper airway CD4 T cells

Memory CD4 T cell phenotypes were examined in NP and MT tissue sites. Upper airway CD8 T_mem_ were noted to be relatively homogeneous, with ∼80-95% of the non-naive CD8 T cells in both MT and NP sites being T_RM_ (Fig. 3A), for both bulk and antigen-specific CD8 T_mem_ (Fig. 3E-I). In contrast, upper airway CD4 T_mem_ displayed greater phenotypic diversity (Fig. 4A). While most non-naive CD4 T cells in both MT and NP tissue sites expressed CD69, only 36% co-expressed CD103 in MT tissues, and only 12% co-expressed CD103 in NP tissues (Fig. 4A). Expanded immunophenotyping across subjects, tissues, and timepoints revealed that MT CD4 T_RM_ most frequently co-expressed CD69 and CXCR6 (Fig. 4A-C). CXCR6 has been shown to be important for nasal CD4 T_RM_ in mice, and associated with protection (*12*). A fraction of both MT (49%) and NP (44%) CD4 T_RM_ co-expressed CD69 and CCR6 (Fig. 4A, D-E). Upper airway CD4 T_mem_ phenotypes were generally stable over time based on surface marker expression (fig. S4A), with expression of CXCR6 and CCR6 notably unchanged over time among MT and NP CD4 T_RM_ (Fig. 4B, D).

**Figure 4.**
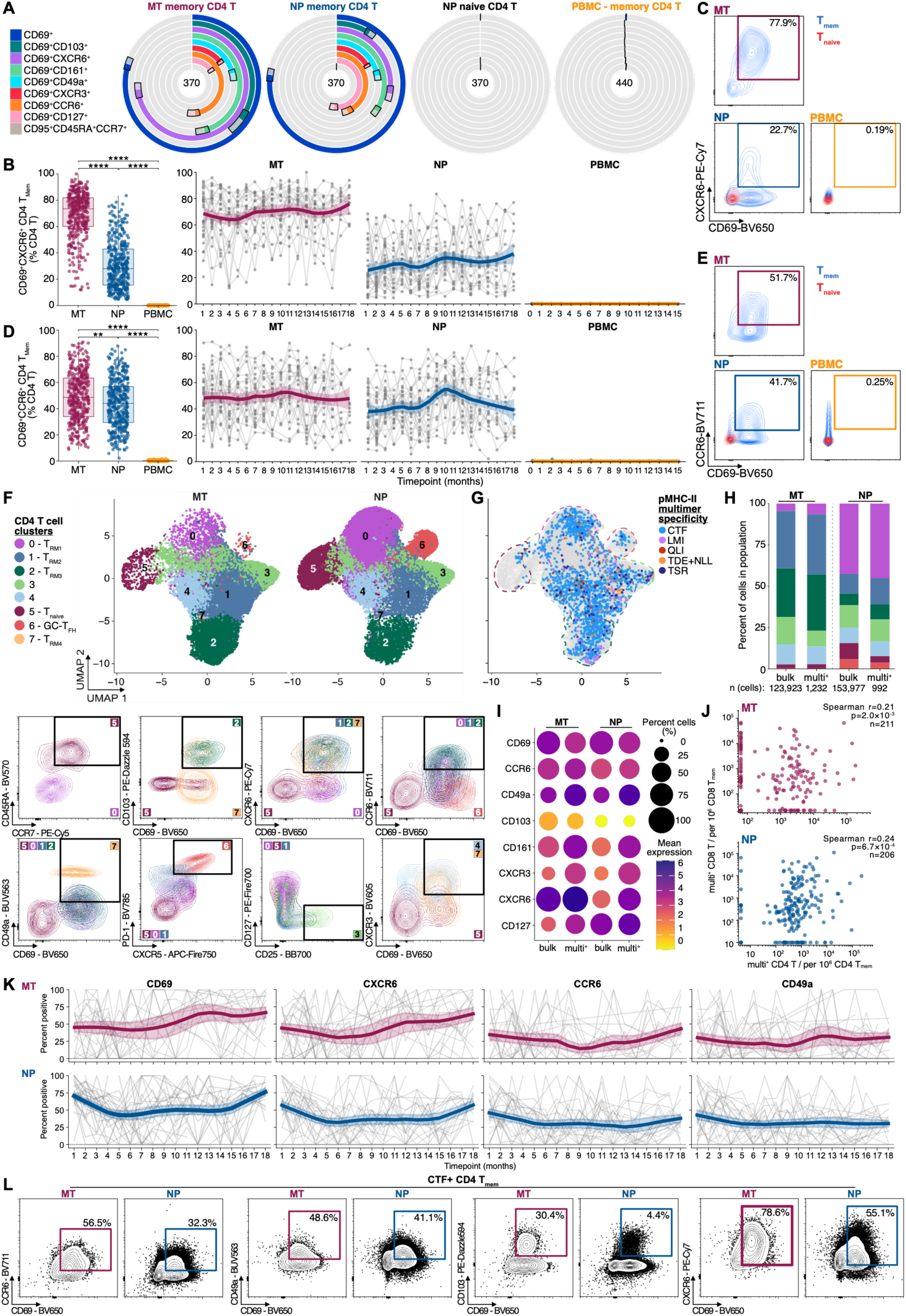
Upper airway CD4 T cell phenotypic features across tissues. A. ‘Belt’ plots with concentric rings demonstrating the proportion of memory CD4 T cells from MT swabs, NP swabs, or PBMC and naive CD4 T cells from NP swabs that express CD69 alone or in combination with other putative T_RM_ markers (values in the centers of the concentric rings = number of samples); medians and 95% CI are shown. B. Cross-sectional (left) and longitudinal (right) co-expression of CD69 and CXCR6 by non-naive CD4 T cells in MT and NP swabs, PBMC. Box and whisker plots display median, IQR, and range. Left: Pairwise Wilcoxon tests with Bonferroni correction for multiple comparison testing; ** = p-value < or = 0.01, **** = p-value < or = 0.0001. Right: Thin gray lines and dots = individual subject’s trajectories and monthly data points. Thick colored lines = group geomean. C. Example flow cytometry contour plots of CD69 and CXCR6 co-expression by non-naive CD4 T cells in MT swabs (black; CD45RA^-^ CCR7^-^, CD45RA^-^CCR7^+^, and CD45RA^+^CCR7^-^), and overlaid naive (red, CD45RA^+^CCR7^+^) versus non-naive (light blue) CD4 T cells in NP swabs and PBMC. % CD69^+^CXCR6^+^ of parental (non-naive CD4 T cells) is shown. D. Cross-sectional (left) and longitudinal (right) co-expression of CD69 and CCR6 by non-naive CD4 T cells in MT and NP swabs, PBMC. Box and whisker plots display median, IQR, and range. Left: Pairwise Wilcoxon tests with Bonferroni correction for multiple comparison testing; ** = p-value < or = 0.01, **** = p-value < or = 0.0001. Right: Thin gray lines and dots = individual subject’s trajectories and monthly data points. Thick colored lines = group geomean. E. Example flow cytometry contour plots of CD69 and CCR6 co-expression by non-naive CD4 T cells in MT swabs (blue, T_Mem_; CD45RA^-^ CCR7^-^, CD45RA^-^CCR7^+^, and CD45RA^+^CCR7^-^). Overlaid naive (red, CD45RA^+^CCR7^+^, T_naive_) versus non-naive (blue, T_Mem_) CD4 T cells in NP swabs and PBMC. % CD69^+^CCR6^+^ of parental (non-naive CD4 T cells) is shown. F. Top: Seurat UMAPs split by anatomic site (MT and NP) of single cell CD8 T cell flow cytometry data from 740 MT and NP samples from 31 subjects. Random downsampling prior to Seurat object construction with up to 5,000 cells per subject and site yielded a Seurat object with 285,228 CD4 T cells. Fourteen surface markers were used as variable features: CD69, CD103, CD49a, CXCR6, CCR6, CD161, CXCR3, CD127, CXCR5, CCR7, CD45RA, PD-1, CD25, and CD95. The result was a total of 8 CD4 T cell clusters at a resolution of 0.2. For visualization, 50,000 cells were randomly subsampled from each object. See Methods and Fig. S4C for additional details. Bottom: Flow cytometry plots generated from the CD4 T cells assigned to each Seurat cluster demonstrating differential marker expression by the CD4 T cell clusters related to naive (CD45RA^+^CCR7^+^) vs memory, T_RM_ (CD69, CD103, CXCR6, CD49a), GC-T_FH_ (CXCR5^+^PD-1^hi^), and other CD4 T cell subset phenotypes. G. Seurat UMAP of the combined downsampled MT and NP data from D as a background of gray, multimer-negative total CD4 T cells with an overlay of multimer-positive CD4 T cells from 25 HLA-matched subjects color-coded by class II multimer(s)-specificity. Cells binding more than one multimer (other than those binding TDE/NLL, where the allele restriction is shared and binding to TDE vs NLL could not be deconvoluted) were excluded. To aid visualization, approximate outlines of the clusters from F are shown in the corresponding colors and SARS2 multimer-positive cells are plotted at an increased point size; n = 2,236 multimer-positive CD4 T cells are shown, of ∼23,000 total pMHC-multimer-positive CD4 T cell events collected. H. Stacked bar plot showing (from left to right) the proportion of bulk MT, multimer-positive MT, bulk NP, and multimer-positive NP CD4 T cells contributing to each of the 8 clusters shown in F; n = total number of bulk and antigen-specific CD4 T cells in the analysis. I. Dot plot with unscaled surface protein expression of CD69 and other putative T_RM_ markers by bulk MT and NP and SARS2 class II multimer-positive MT and NP CD4 T cells. Dot size = % of total CD4 T cells expressing CD69 or % of total CD69^+^ CD4 T cells expressing each additional marker, per sample; averaged across samples per site for bulk cells (MT ∼450,000 and NP ∼11 million CD4 T cells from flow cytometry data), or averaged first across epitope specificities per subject per timepoint then across subjects per site for multimer-positive cells (MT 2,026 and NP 19,718 total pMHC-II multimer-positive CD4 T cells), with a minimum of 3 CD69^+^ cells per sample required. Dot color = mean asinh fluorescence intensity (cofactor 150, fixed color scale 0–6, dark = high expression) computed from the projected multimer-positive Seurat object cell subset (∼23,000 CD4 T cells) or from the Seurat object for bulk CD4 T cells. J. Correlations between SARS2-specific memory CD8 and CD4 T cell frequencies for all SARS2 class I and II multimers (averaged by the number of pMHC-I/II multimers per sample) for matched MT (top, red) and NP (bottom, blue) swab samples (n = 111) from the same subjects and time points. Zero values were set to the minimum threshold for positivity for SARS2 class I/II multimers by site (MT vs NP). K. T_RM_ marker (CD69, CXCR6, CCR6, CD49a) expression stability in SARS2 class II multimer-positive CD4 T_mem_ over time by flow cytometry. % of pooled multimer-positive CD4 T_mem_ in MT (n = 219) or NP (n = 315) swabs expressing each indicated marker. Data from swabs collected monthly for up to 18 months from 29-30 subjects (MT = 29, NP = 30; individual timepoints include 6–25 subjects). Thin gray lines represent individual subject’s trajectories. Thick colored line = LOWESS-smoothed mean; shaded band indicates ±SEM. L. Example concatenated flow cytometry data of T_RM_ marker (CD69 and CCR6, CD49a, CD103, CXCR6) co-expression by SARS2 CTF pMHC-II positive memory CD4 T cells in n = 231 MT and 231 NP swabs.

Unsupervised analysis of high-parameter spectral flow cytometry data was performed to further characterize upper airway CD4 T cells. Multiple CD4 T_RM_ and other CD4 T cell subsets were identified from MT and NP tissue sites, with site-specific differences in populations, including an enrichment of naive CD4 T cells and germinal center T follicular helper cells (GC-T_FH_) in NP swabs (Fig. 4F, H; fig. S4B-C). SARS2 pMHC-II multimer-specific cells were distributed across memory CD4 and T_FH_ clusters and primarily found in clusters with phenotypes consistent with CD4 T_RM_ (Fig. 4G-H). SARS2-specific and bulk CD4 T_RM_ were compared, from both MT and NP tissue sites. While only 41% of bulk NP CD4 T_RM_ expressed CXCR6, a substantially higher proportion of SARS2-specific CD4 T_RM_ expressed CXCR6 (73%) (Fig. 4I). SARS2-specific MT and NP CD4 T_RM_ also more frequently expressed CD49a and CXCR3 than bulk upper airway CD4 T_RM_ (Fig. 4I).

Relationships between memory CD8 and CD4 T_mem_ abundance in the upper airway were examined. SARS2-specific CD8 and CD4 T cell frequencies were positively associated in the MT epithelium (r = 0.21, p = 0.002. Fig. 4J). SARS2-specific CD8 and CD4 T cell frequencies were also positively correlated in NP samples (r = 0.24, p = 6.7 x 10^-4^. Fig. 4J).

SARS2-specific upper airway memory CD8 T cells exhibited stable phenotypes over time in the tissue (Fig. 3I). We therefore examined the stability of SARS2-specific upper airway memory CD4 T cell phenotypes over time. SARS2-specific MT and NP CD4 T_mem_ displayed relatively stable T_RM_ phenotypes over a period of 18 months (Fig. 4K-L, fig. S4A). Influenza A multimer-specific CD4 T_mem_ possessed similar T_RM_ phenotypes in both human MT and NP tissue sites (fig. S4D). Thus, antigen-specific memory CD8 and CD4 T cells in upper airway epithelial mucosa and lymphoid tissue are positively correlated, these cells persist over time, and the persistence of these cells is associated with a T_RM_ program.

## Discussion

These data show direct evidence for long-lived, antigen-specific human upper airway CD8 and CD4 T_RM_. Employing minimally invasive longitudinal swab sampling of upper respiratory tract tissues and an array of pMHC multimers, we were able to track rare, respiratory virus-specific CD8 and CD4 T cells for > 18 months with epitope level specificity. Circulating CD8 and CD4 memory T cells with the same specificities were also tracked for at least 12 months. Memory T cell frequencies were stably maintained, without an apparent half-life, across NP, blood, and mucosal epithelial tissues.

Epitope-specific SARS2-specific memory CD8 T cells and CD4 T cells were commonly detected in upper airway tissue sites in this cohort. By utilizing a total of 13 SARS2 pMHC multimers (7 class I and 6 class II), T cell memory to SARS2 was assessed at an HLA coverage level indicating that 90% of the global population would be expected to have detectable memory T cells in upper airway tissue to at least one of the epitopes tested (2.8 epitopes on average) (*33*).

As suggested by prior TCR sequencing work (*8*), there is overlap between circulating and upper airway T cell specificity for SARS2 antigens. There is also overlap between circulating and upper airway T cell specificity for immunodominant Influenza A antigens (Fig 1 & 2), consistent with clonotype sharing observed between blood and lung (*4*, *32*). Despite shared pMHC specificity, antigen-specific circulating and upper airway memory T cells were phenotypically distinct. SARS2 and Influenza A-specific CD8 and CD4 T cells in both mucosal epithelium and secondary lymphoid upper airway tissues demonstrated stable T_RM_ phenotypes with features pointing to a conserved program underlying maintenance of these T cells within the upper respiratory tract. While CD8 T cells were phenotypically similar regardless of site (MT or NP), CD4 T cells displayed greater phenotypic diversity, indicating more memory CD4 T cell subset specialization in upper respiratory tract tissue niches.

Several symptomatic SARS2 infections were reported or detected during the study, and upper airway antigen-specific T cell frequencies were sensitive for the detection of infection events. Overall, the data indicate that CD8 and CD4 T_RM_ present in the human upper respiratory tract are poised to surveil and rapidly respond to respiratory pathogens.

Limitations of this study include that participants in this study were generally healthy adults, and thus the results may not be generalizable to the extremes of age (*34*, *35*). Given that SARS2 is endemic and Influenza A circulates seasonally, antigen exposure cannot be excluded. Mitigating this limitation, this cohort was sampled monthly and examined by multiple approaches for evidence of infections. Local transmission of SARS2 was relatively low during this period (*36*). Lastly, this study examined individuals for 18 months; while that is the longest to date, studying even longer time periods for longitudinal immune memory would be of value.

Respiratory infections remain a major global public health burden. The data here indicate that mucosal immune memory in humans may offer an additional layer of protection against respiratory pathogens. Major efforts are underway worldwide to develop next generation vaccines, with an emphasis on eliciting and enhancing mucosal immunity (*5*, *18*, *37–40*). The findings herein demonstrate that durable antigen-specific CD8 T_RM_ and CD4 T_RM_ maintenance in human mucosal epithelial tissue, and T_RM_ in mucosal lymphoid tissue, is possible, and that next generation vaccines may be able to leverage mucosal T cells to generate protective immune memory.

## Acknowledgments

We would like to thank the participants for providing the samples and clinical information that made this study possible. We are grateful to the LJI Clinical Studies Core staff, in particular Gina Levi, Jasmine Cardenas, and Noemi Chavez for collecting most of the samples used in this study and providing clinical coordination services. We thank the NIH Tetramer Core Facility (NIH Contract 75N93020D00005 and RRID:SCR_026557) for providing SARS2 and influenza A reagents (see Table S3 for details). The blood drop, epithelium, and adenoid icons shown in Fig. 1 and 2 were generated with the assistance of Google Gemini 3.1 Pro. Fig. S1A was generated in BioRender. R scripts for data analysis and figure generation in R were written with assistance from Claude Sonnet 4.6-5 (Anthropic).

## Funding

This work was supported in part by the National Institute of Allergy and Infectious Diseases (NIAID) of the National Institutes of Health (NIH), Department of Health and Human Services, under award AI142742 Collaborative Center for Human Immunology (S.C.) and award K08AI196260 (S.R.).

## Author contributions

HLA-typing and other clinical data for recruitment were provided by A.S. with data curation provided by A.A., and A.F. E.J.P. and S.M. performed HLA-typing. A.S. helped with calculating projected population coverage for pMHC reagents. T.D. and M.K.J. designed, generated, and provided the SARS2 spike pMHC-II CTF monomer and tetramer, and provided guidance on conditions for multimerization and multimer staining. S.I.R., F.F., and S.C. designed the experiments. S.I.R., F.F., and P.G.L. performed nasal swab sample processing and flow cytometry. N.H., L.G.-O., O.E., and P.G.L. performed blood processing and carried out serological assays (MSD V-PLEX, enzyme-linked immunosorbent assays). S.I.R. and F.F. contributed to flow cytometry, serology, and other data analyses. S.I.R. and F.F. generated the figures. S.I.R., F.F., and S.C. wrote the original manuscript draft. All authors reviewed and edited the manuscript. Supervision and funding were provided by S.R. and S.C.

## Competing Interests

The authors have no competing interests to declare.

## Data and Materials Availability

All the data needed to evaluate the conclusions in the paper are present in the main text, supplementary materials, and source data. All materials are available through commercial sources or the NIH Tetramer Core facility.

## Supplementary Materials

Figure S1-S4

Figure S1 related to Figure 1

Figure S2 related to Figure 2

Figure S3 related to Figure 3

Figure S4 related to Figure 4

Tables S1-S5

Table S1 study cohort demographics and clinical history

Table S2 pertinent HLA allele information for the study cohort

Table S3 SARS2 and influenza A pMHC-I/II multimer list

Table S4 List of flow cytometry antibodies in panel v2

Table S5 List of flow cytometry antibodies in panel v3

Source Data S1-S2

Data S1 data related to all main text and supplementary figures other than single cell analysis

Data S2 data related to high-dimensional flow cytometry single cell analyses

## Notes

### Competing Interest Statement

The authors have declared no competing interest.

## References

1. S. Crotty, Immunological memory to vaccines. Immunity 59, 813–832 (2026).

2. M. M. L. Poon, D. P. Caron, Z. Wang, S. B. Wells, D. Chen, W. Meng, P. A. Szabo, N. Lam, M. Kubota, R. Matsumoto, A. Rahman, E. T. Luning Prak, Y. Shen, P. A. Sims, D. L. Farber, Tissue adaptation and clonal segregation of human memory T cells in barrier sites. Nat Immunol 24, 309–319 (2023).

3. B. V. Kumar, W. Ma, M. Miron, T. Granot, R. S. Guyer, D. J. Carpenter, T. Senda, X. Sun, S.-H. Ho, H. Lerner, A. L. Friedman, Y. Shen, D. L. Farber, Human Tissue-Resident Memory T Cells Are Defined by Core Transcriptional and Functional Signatures in Lymphoid and Mucosal Sites. Cell Rep 20, 2921–2934 (2017).

4. V. Fajardo-Rosas, A. Ehtram, I. Marchand-Casas, S. J. Chee, M. Mondal, C. Ferrández Alaminos, F. E. Castañeda-Castro, Z. Roy, L. Chudley, M. Shackcloth, J. Cave, A. Alzetani, E. Woo, E. Phillips, S. Mallal, A. Grifoni, B. J. Schmiedel, A. Sette, B. Peters, C. H. Ottensmeier, P. Vijayanand, Human lungs maintain tissue-resident memory T cells against a broad spectrum of pathogens. Nat Immunol 27, 1749–1761 (2026).

5. J. Rosenheim, B. Bender, J. Gilmour, K. Jambo, J. U. S. Jensen, D. King, H. Kløverpris, S. Taylor, M. Boaz, C. Chiu, S. Crotty, C. Dahlke, K. Dheda, S. Fortune, S. L. Higham, M. Holm, S. P. Jochems, F. Krammer, C. S. Lindestam Arlehamn, H. W. Mankouri, H. V. Marquart, H. McShane, R. Mortensen, E. Nemes, J. Ordovas-Montanes, P. C. Roberts, M. Ruhwald, K. L. Schully, C. Thålin, R. S. Thwaites, T. Y. Wang, H. B. Juel, Conference report: airway mucosal sampling and immune analysis. Vaccine 88, 128944 (2026).

6. J. Davis-Porada, A. B. George, N. Lam, D. P. Caron, J. I. Gray, J. Huang, J. Hwu, S. B. Wells, R. Matsumoto, M. Kubota, Y. Lee, R. Morrison-Colvin, I. J. Jensen, B. B. Ural, N. Shaabani, D. Weiskopf, A. Grifoni, A. Sette, P. A. Szabo, J. R. Teijaro, P. A. Sims, D. L. Farber, Maintenance and functional regulation of immune memory to COVID-19 vaccines in tissues. Immunity 57, 2895–2913.e8 (2024).

7. N. Lam, Y. Lee, D. L. Farber, A guide to adaptive immune memory. Nat Rev Immunol 24, 810–829 (2024).

8. S. I. Ramirez, F. Faraji, L. B. Hills, P. G. Lopez, B. Goodwin, H. D. Stacey, H. J. Sutton, K. M. Hastie, E. O. Saphire, H. J. Kim, S. Mashoof, C. H. Yan, A. S. DeConde, G. Levi, S. Crotty, Immunological memory diversity in the human upper airway. Nature 632, 630–636 (2024).

9. S. I. Ramirez, P. G. Lopez, F. Faraji, U. M. Parikh, A. Heaps, J. Ritz, C. Moser, J. J. Eron, D. Wohl, J. Currier, E. S. Daar, A. Greninger, P. Klekotka, A. Grifoni, D. Weiskopf, A. Sette, B. Peters, M. D. Hughes, K. W. Chew, D. M. Smith, S. Crotty, Accelerating COVID-19 Therapeutic Interventions and Vaccines-2 (ACTIV-2)/A5401 Study Team, Early antiviral CD4+ and CD8+ T cells are associated with upper airway clearance of SARS-CoV-2. JCI Insight 9, e186078 (2024).

10. H. R. Wagstaffe, R. S. Thwaites, J. K. Sidhu, R. G. H. Lindeboom, L. Kretschmer, K. B. Worlock, L. M. Dratva, A. Huang, S. Ascough, L. Papargyris, R. McKendry, A. M. Collins, J. Xu, N.-M. Lemm, B. Killingley, M. Kalinova, A. Mann, A. Catchpole, L. Swadling, J. S. Tsang, M. K. Maini, M. Noursadeghi, M. Z. Nikolić, S. A. Teichmann, P. J. M. Openshaw, C. Chiu, Pre-existing and early cellular immune factors correlate with functionally complete protection against primary controlled human SARS-CoV-2 infection. Nat Commun 17, 312 (2025).

11. A. Pizzolla, T. H. O. Nguyen, J. M. Smith, A. G. Brooks, K. Kedzieska, W. R. Heath, P. C. Reading, L. M. Wakim, Resident memory CD8+ T cells in the upper respiratory tract prevent pulmonary influenza virus infection. Sci Immunol 2, eaam6970 (2017).

12. N. R. Mathew, R. Gailleton, L. Scharf, K. Schön, J. Enriquez, H. Axelsson, A. Strömberg, N. Lycke, M. Bemark, K.-W. Tang, D. Angeletti, Nasal CD4+ tissue-resident memory T cells provide cross-protective immunity to influenza. J Exp Med 223, e20251793 (2026).

13. S. W. Kazer, J. M. L. Walsh, L. J. Juttukonda, J. Ordovas-Montanes, Nasal immunity in respiratory viral infection, transmission, and protection. Immunity, S1074-7613(26)00262–1 (2026).

14. J. I. Gray, D. L. Farber, Tissue-Resident Immune Cells in Humans. Annu Rev Immunol 40, 195–220 (2022).

15. A. Grifoni, A. Sette, From Alpha to omicron: The response of T cells. Curr Res Immunol 3, 146–150 (2022).

16. V. Joag, B. N. Bimber, C. F. Quarnstrom, V. S. Bollimpelli, J. M. Schenkel, K. A. Fraser, M. Bertogliat, A. G. Soerens, J. M. Stolley, S. D. O’Flanagan, P. C. Rosato, N. V. Gavil, M. Künzli, J. S. Mitchell, T. Legere, S. Jean, A. A. Upadhyay, C. Y. Kang, J. Gibbs, J. W. Yewdell, B. T. Fife, H. Park, S. G. Hansen, S. E. Bosinger, G. N. Barber, P. J. Skinner, V. Vezys, E. Hunter, L. J. Picker, R. R. Amara, D. Masopust, Primate resident memory T cells activate humoral and stromal immunity. Immunity 58, 2541–2555.e6 (2025).

17. R. G. H. Lindeboom, K. B. Worlock, L. M. Dratva, M. Yoshida, D. Scobie, H. R. Wagstaffe, L. Richardson, A. Wilbrey-Clark, J. L. Barnes, L. Kretschmer, K. Polanski, J. Allen-Hyttinen, P. Mehta, D. Sumanaweera, J. M. Boccacino, W. Sungnak, R. Elmentaite, N. Huang, L. Mamanova, R. Kapuge, L. Bolt, E. Prigmore, B. Killingley, M. Kalinova, M. Mayer, A. Boyers, A. Mann, L. Swadling, M. N. J. Woodall, S. Ellis, C. M. Smith, V. H. Teixeira, S. M. Janes, R. C. Chambers, M. Haniffa, A. Catchpole, R. Heyderman, M. Noursadeghi, B. Chain, A. Mayer, K. B. Meyer, C. Chiu, M. Z. Nikolić, S. A. Teichmann, Human SARS-CoV-2 challenge uncovers local and systemic response dynamics. Nature 631, 189–198 (2024).

18. T. N. Burn, L. K. Mackay, Spatial organization of tissue-resident memory T cells. Immunity 59, 863–877 (2026).

19. M. Heeg, A. W. Goldrath, Insights into phenotypic and functional CD8+ TRM heterogeneity. Immunol Rev 316, 8–22 (2023).

20. A. Tarke, C. H. Coelho, Z. Zhang, J. M. Dan, E. D. Yu, N. Methot, N. I. Bloom, B. Goodwin, E. Phillips, S. Mallal, J. Sidney, G. Filaci, D. Weiskopf, R. da Silva Antunes, S. Crotty, A. Grifoni, A. Sette, SARS-CoV-2 vaccination induces immunological T cell memory able to cross-recognize variants from Alpha to Omicron. Cell 185, 847–859.e11 (2022).

21. A. Zhu, Z. Chen, Q. Yan, M. Jiang, X. Liu, Z. Li, N. Li, C. Tang, W. Jian, J. He, L. Chen, J. Cheng, C. Chen, T. Tang, Z. Xu, Q. Hu, F. Li, Y. Wang, J. Sun, Z. Zhuang, L. Wen, J. Zhuo, D. Liu, Y. Zhang, X. Huang, S. Li, Q. Zeng, F. Chen, L. Zhou, D. Liu, C. Zhong, Y. Chen, S. Li, K. Liang, N. Zhong, X. Zhang, J. Chen, X. Chen, Y. Xu, N. Zhong, J. Zhao, J. Zhao, Robust mucosal SARS-CoV-2-specific T cells effectively combat COVID-19 and establish polyfunctional resident memory in patient lungs. Nat Immunol 26, 459–472 (2025).

22. D. Masopust, A. G. Soerens, Tissue-Resident T Cells and Other Resident Leukocytes. Annu Rev Immunol 37, 521–546 (2019).

23. D. Goldblatt, G. Alter, S. Crotty, S. A. Plotkin, Correlates of protection against SARS-CoV-2 infection and COVID-19 disease. Immunol Rev 310, 6–26 (2022).

24. J. M. E. Lim, A. T. Tan, N. Le Bert, S. K. Hang, J. G. H. Low, A. Bertoletti, SARS-CoV-2 breakthrough infection in vaccinees induces virus-specific nasal-resident CD8+ and CD4+ T cells of broad specificity. J Exp Med 219, e20220780 (2022).

25. R. S. Akondy, M. Fitch, S. Edupuganti, S. Yang, H. T. Kissick, K. W. Li, B. A. Youngblood, H. A. Abdelsamed, D. J. McGuire, K. W. Cohen, G. Alexe, S. Nagar, M. M. McCausland, S. Gupta, P. Tata, W. N. Haining, M. J. McElrath, D. Zhang, B. Hu, W. J. Greenleaf, J. J. Goronzy, M. J. Mulligan, M. Hellerstein, R. Ahmed, Origin and differentiation of human memory CD8 T cells after vaccination. Nature 552, 362–367 (2017).

26. A. A. Minervina, M. V. Pogorelyy, A. M. Kirk, J. C. Crawford, E. K. Allen, C.-H. Chou, R. C. Mettelman, K. J. Allison, C.-Y. Lin, D. C. Brice, X. Zhu, K. Vegesana, G. Wu, S. Trivedi, P. Kottapalli, D. Darnell, S. McNeely, S. R. Olsen, S. Schultz-Cherry, J. H. Estepp, SJTRC Study Team, M. A. McGargill, J. Wolf, P. G. Thomas, SARS-CoV-2 antigen exposure history shapes phenotypes and specificity of memory CD8+ T cells. Nat Immunol 23, 781–790 (2022).

27. M. Koutsakos, W. S. Lee, A. Reynaldi, H.-X. Tan, G. Gare, P. Kinsella, K. C. Liew, G. Taiaroa, D. A. Williamson, H. E. Kent, E. Stadler, D. Cromer, D. S. Khoury, A. K. Wheatley, J. A. Juno, M. P. Davenport, S. J. Kent, The magnitude and timing of recalled immunity after breakthrough infection is shaped by SARS-CoV-2 variants. Immunity 55, 1316–1326.e4 (2022).

28. T. H. O. Nguyen, L. C. Rowntree, J. Petersen, B. Y. Chua, L. Hensen, L. Kedzierski, C. E. van de Sandt, P. Chaurasia, H.-X. Tan, J. R. Habel, W. Zhang, L. F. Allen, L. Earnest, K. Y. Mak, J. A. Juno, K. Wragg, F. L. Mordant, F. Amanat, F. Krammer, N. A. Mifsud, D. L. Doolan, K. L. Flanagan, S. Sonda, J. Kaur, L. M. Wakim, G. P. Westall, F. James, E. Mouhtouris, C. L. Gordon, N. E. Holmes, O. C. Smibert, J. A. Trubiano, A. C. Cheng, P. Harcourt, P. Clifton, J. C. Crawford, P. G. Thomas, A. K. Wheatley, S. J. Kent, J. Rossjohn, J. Torresi, K. Kedzierska, CD8+ T cells specific for an immunodominant SARS-CoV-2 nucleocapsid epitope display high naive precursor frequency and TCR promiscuity. Immunity 54, 1066–1082.e5 (2021).

29. K. M. Wragg, W. S. Lee, M. Koutsakos, H.-X. Tan, T. Amarasena, A. Reynaldi, G. Gare, P. Konstandopoulos, K. R. Field, R. Esterbauer, H. E. Kent, M. P. Davenport, A. K. Wheatley, S. J. Kent, J. A. Juno, Establishment and recall of SARS-CoV-2 spike epitope-specific CD4+ T cell memory. Nat Immunol 23, 768–780 (2022).

30. N. Borcherding, W. Kim, M. Quinn, F. Han, J. Q. Zhou, A. J. Sturtz, A. J. Schmitz, T. Lei, S. A. Schattgen, M. K. Klebert, T. Suessen, W. D. Middleton, C. W. Goss, C. Liu, J. C. Crawford, P. G. Thomas, S. A. Teefey, R. M. Presti, J. A. O’Halloran, J. S. Turner, A. H. Ellebedy, P. A. Mudd, CD4+ T cells exhibit distinct transcriptional phenotypes in the lymph nodes and blood following mRNA vaccination in humans. Nat Immunol 25, 1731–1741 (2024).

31. P. A. Mudd, A. A. Minervina, M. V. Pogorelyy, J. S. Turner, W. Kim, E. Kalaidina, J. Petersen, A. J. Schmitz, T. Lei, A. Haile, A. M. Kirk, R. C. Mettelman, J. C. Crawford, T. H. O. Nguyen, L. C. Rowntree, E. Rosati, K. A. Richards, A. J. Sant, M. K. Klebert, T. Suessen, W. D. Middleton, SJTRC Study Team, J. Wolf, S. A. Teefey, J. A. O’Halloran, R. M. Presti, K. Kedzierska, J. Rossjohn, P. G. Thomas, A. H. Ellebedy, SARS-CoV-2 mRNA vaccination elicits a robust and persistent T follicular helper cell response in humans. Cell 185, 603–613.e15 (2022).

32. S. Sant, L. Grzelak, Z. Wang, A. Pizzolla, M. Koutsakos, J. Crowe, T. Loudovaris, S. I. Mannering, G. P. Westall, L. M. Wakim, J. Rossjohn, S. Gras, M. Richards, J. Xu, P. G. Thomas, L. Loh, T. H. O. Nguyen, K. Kedzierska, Single-Cell Approach to Influenza-Specific CD8+ T Cell Receptor Repertoires Across Different Age Groups, Tissues, and Following Influenza Virus Infection. Front Immunol 9, 1453 (2018).

33. H.-H. Bui, J. Sidney, K. Dinh, S. Southwood, M. J. Newman, A. Sette, Predicting population coverage of T-cell epitope-based diagnostics and vaccines. BMC Bioinformatics 7, 153 (2006).

34. P. A. Szabo, H. M. Levitin, S. Nargund, T. J. Connors, D. Chen, J. Jin, P. Thapa, R. Guyer, D. P. Caron, J. I. Gray, R. Matsumoto, M. Kubota, M. Brusko, T. M. Brusko, D. L. Farber, P. A. Sims, Distinct transcription factors control tissue adaptation and effector function in infant and adult memory T cells. Nat Immunol 27, 1517–1527 (2026).

35. S. Adamo, J. G. Rurik, C. E. Gustafson, M. Buggert, Memory T cell aging and rejuvenation. Immunity 59, 878–896 (2026).

36. County of San Diego Wastewater Surveillance Dashboard; https://www.sandiegocounty.gov/content/sdc/hhsa/programs/phs/phs_laboratory/WastewaterDashboard.html.

37. G. Neumann, B. J. Cowling, H. Chen, S. Stertz, B. Manicassamy, I. G. Barr, T. M. Uyeki, Y. Kawaoka, Influenza. Nat Rev Dis Primers 12, 51 (2026).

38. R. C. Mettelman, E. K. Allen, P. G. Thomas, Mucosal immune responses to infection and vaccination in the respiratory tract. Immunity 55, 749–780 (2022).

39. T. Inoue, W. Ise, T. Kurosaki, GC B Cells, Tfh Cells, and Influenza bnAbs: Insights for Improving Universal Vaccine Design. Immunol Rev 341, e70153 (2026).

40. D.-I. Kwon, S. H. Bhagchandani, S. A. Ehrenzeller, A. Iwasaki, Harnessing mucosal immunity for protective vaccines. Nat Rev Immunol 26, 507–524 (2026).

41. A. Tarke, J. Sidney, C. K. Kidd, J. M. Dan, S. I. Ramirez, E. D. Yu, J. Mateus, R. da Silva Antunes, E. Moore, P. Rubiro, N. Methot, E. Phillips, S. Mallal, A. Frazier, S. A. Rawlings, J. A. Greenbaum, B. Peters, D. M. Smith, S. Crotty, D. Weiskopf, A. Grifoni, A. Sette, Comprehensive analysis of T cell immunodominance and immunoprevalence of SARS-CoV-2 epitopes in COVID-19 cases. Cell Rep Med 2, 100204 (2021).

42. Y. Hao, T. Stuart, M. H. Kowalski, S. Choudhary, P. Hoffman, A. Hartman, A. Srivastava, G. Molla, S. Madad, C. Fernandez-Granda, R. Satija, Dictionary learning for integrative, multimodal and scalable single-cell analysis. Nat Biotechnol 42, 293–304 (2024).

43. Q. Xu, L. Shi, B. L. P. Dizon, R. J. Beam, P. Mudd, P. L. Schwartzberg, K. Manthiram, Efficient Tonsillar T Follicular Helper Cell Processing and Functional Analysis through High-dimensional Flow Cytometry. J Vis Exp, doi: 10.3791/67188 (2025).

