## Supplementary materials for "Durability and diversity of human upper airway T cell memory"

**The PDF file includes:**

Materials and Methods
Figs. S1 to S4
Tables S1 to S5

**Other Supplementary Materials for this manuscript include the following:**

Data S1 to S2
MDAR Reproducibility Checklist

### Materials and Methods

Human Subjects

Adult subjects ages 18-70 years old were enrolled under the human subjects study protocol #VD-275 approved by the La Jolla Institute for Immunology (LJI) Institutional Review Board. Subjects were recruited based on the availability and relevance of HLA-typing results generated by Next Generation Sequencing (NGS) and enrolled in the study on a rolling basis. Subjects were enrolled between June 2024 and January 2025. All subjects provided written consent for study participation and verbal consent for sample collections.

Sample Collections

Nasal swabs were collected from two distinct sites to sample mucosal and lymphatic tissues present within the upper airway. Nasal mid-inferior turbinate (MT) and nasopharyngeal (NP) swab sample collections were performed by a limited number of LJI Clinical Core or trained study staff using universal precautions and standard collection protocols, as previously described (*8*). Briefly, a Puritan Hydraflock swab (cat # 25-3306-H) was gently inserted into one nostril and passed along the floor of the nasal cavity to reach the posterior nasopharynx (~7-8 cm) in an attempt to sample adenoid tissue (NP swab), or approximately half this distance (~3-4 cm) to sample the inferior nasal turbinate mucosa (MT swabs) at a shallower depth that would ensure that the nasopharynx was not being sampled. Once the swab reached the desired sampling depth within the nasal cavity, the swab was rotated clockwise for 5-10 full rotations (as tolerated) while maintaining pressure against the posterior wall of the nasopharynx (NP swabs) or against the inferior turbinate mucosa (MT swabs) to collect cells from the respective anatomic site. Swab sampling was performed bilaterally as tolerated, unless an anatomic barrier precluded bilateral swab sampling. MT swabs were collected first to avoid the possibility of 'cross-contamination' with adenoid/NP cells. Peripheral blood samples were also collected by LJI Clinical Core staff using universal precautions and standard phlebotomy protocols. Samples were collected between late June 2024 and late June 2026.

Sample Processing

All samples were processed on the same day as collection using standard laboratory protocols, as previously described (*8*). Briefly, whole peripheral blood was collected in EDTA tubes (BD) and stored at room temperature until processed to isolate plasma and peripheral blood mononuclear cells (PBMC). Plasma and PBMC were aliquoted and cryopreserved for future use. PBMC were later thawed for staining using standard methods. After collection, nasal swabs were placed in 15 mL conical collection tubes containing 2 mL of complete cell culture medium (RPMI 1640 (Corning) supplemented with 1% Penicillin/Streptomycin (Gibco), 1% GlutaMAX (Gibco), and 5% FBS) and samples were stored at 4° Celsius until processed. Upon processing, collection tubes containing the swabs were pulse vortexed at medium to high speed to release cells from the swabs. The swab tip was then placed in a clean 40 μm filter resting on top of a 50 mL conical tube and rinsed with the 2 mL of collection medium and then another 10 mL of fresh cell culture medium. The filtered cell suspension was centrifuged in 20°C centrifuge at 500 rcf for 7 minutes. The supernatant was discarded, and the cell pellet was resuspended in 100 μL of room temperature complete cell culture medium to undergo multimer staining.

Quality control (QC) metrics for PBMC and nasal swab samples

Upon thawing, PBMC samples were subjected to cell counts and viability staining (Cytek Muse), and a lymphocyte viability threshold of > 85% viability after thawing was applied to all PBMC samples. Given individual variation between subjects in terms of the frequencies of total and memory CD8 and CD4 T cells in MT and NP swabs and inherent sampling variability, T cell count based metrics were applied to nasal swab samples. For NP swabs, samples with no detectable multimer responses and fewer than 1,000 CD8 or CD4 T cells were excluded from class I or class II multimer analyses, respectively. For MT swabs, samples with no detectable multimer responses and fewer than 100 CD8 or CD4 T cells were excluded from multimer analyses. Samples that had detectable multimer responses but did not meet the CD8 or CD4 T cell anatomic site-specific QC threshold were plotted on longitudinal multimer plots (solid black circles) and counted as positive responses but not included in group geomean calculations. For inclusion in longitudinal group plots of MT, NP, or PBMC multimer-specific responses, each subject was required to have true positive responses (QC criteria met and positive response observed) for the given multimer across three or more sampling time points with at least two appropriate multimer-positive (CD8 or CD4) T cells per time point.

Adenoid tissue in the posterior nasopharynx was the target of NP swab sampling. Additional QC metrics beyond those described above for NP swab sample inclusion in analyses were applied to define successful adenoid sampling. Successful adenoid sampling was ultimately determined by a total B cell count threshold--NP swabs with 1,548 or more total B cells were considered adenoid NP swabs (total B cell count threshold for the top 5% of MT swabs). NP swabs that failed to meet the total B cell metric for adenoid swabs were considered non-adenoid NP swabs. Additional per-swab cell count metrics were examined to assess germinal center B cell and GC-$T_{FH}$ sampling by NP swabs (fig. S2M).

Antigen exposure reporting and fold-change analyses

This was an observational, non-interventional study. Participants were asked to report any new upper respiratory infections or symptoms and new vaccinations at each study visit. When respiratory infections were reported, participants were asked if diagnostic testing was performed and whether they tested positive for SARS2 or influenza infections. The timing of sample collections was not coordinated with the timing of reported symptoms or Ag exposure events. Additionally, participants were not excluded from the study based on reported symptoms or Ag exposures. In total, 23 participants underwent longitudinal blood draws every ~3 months and monthly nasal swab sampling over the course of 12 months or greater and had samples that were included in Ag exposure analyses.

Fold-change (FC) values for multimer responses were calculated as the post-event frequency divided by the pre-event frequency of pMHC-I/II multimer positive cells per million non-naive CD8/CD4 T cells for each anatomic site (MT, NP, PBMC). FC values were also calculated for SARS2-specific IgG as the post-event IgG titer divided by the pre-event IgG titer. Data from the nasal swab and blood samples collected closest to the reported BTI or vaccination event were used for FC calculations. Only samples collected within 90-days of the reported event were considered. For suspected events, any paired (sequential) samples collected within the suspected event period could qualify. The proportion of four-fold or greater multimer increases were calculated for reported BTI/suspected infections, reported vaccinations, non-exposure event time points. Fisher's exact testing was performed to compare the proportion of reported/suspected events versus the proportion of all "other" non-event with FC > or = 4. Benjamini-Hochberg correction was applied for multiple comparisons. FC comparisons were also made between the % of multimer-positive or bulk memory CD8 or CD4 T cells in MT, NP, or PBMC samples expressing T cell activation markers for reported and suspected events. Unlike multimer frequencies, there was no statistically significant pattern of Ag-specific or bulk T cell activation marker upregulation post-exposure by Fisher's exact testing with corrections for multiple comparisons.

Serology

SARS2-specific N and S IgG serology data was generated for the 23 subjects who provided longitudinal blood (plasma) samples over the course of ~12 months. Pre-pandemic plasma and pooled plasma from unrelated subjects with hybrid immunity (SARS2 infection plus COVID vaccination) were included as negative and positive controls, respectively. IgG data for primary serologic analysis was generated using the V-Plex SARS-CoV-2 Panel 39 Kit (K15738U-4) and Meso Scale Discovery (MSD) MULTI-SPOT Assay System, in accordance with the manufacturer's protocol and recommendations. Briefly, plates and diluents were equilibrated to room temperature, and all plasma samples and reference controls were thawed on ice prior to performing the assay. MSD Blocker A Kit (R93AA-2) was used to make blocking buffer. Plates were blocked for 30 minutes at room temperature on a shaker set to 700 rpm. Reference Standard 2 was serially diluted in Diluent 100 (MSD, R50AA-3). Once thawed, plasma samples were equilibrated to room temperature then diluted either 1:5,000 or 1:50,000 in Diluent 100. Plates were washed three times with 1x wash buffer. Each dilution of Reference Standard 2 (technical replicates), Diluent 100 (blank), Serology Control 2.1 from the Serology Control Pack 2 (C4731-1), and the diluted experimental samples were then plated and incubated for two hours at room temperature on a shaker set to 700 rpm. After this incubation, the plates were washed three times with 1x wash buffer. Detection antibody (MSD GOLD SULFO-TAG Anti-Human IgG Antibody, D21ADF3) was prepared in Diluent 100, then added to the plates, and incubated at room temperature for one hour on a shaker set to 700 rpm. After the incubation, the plates were washed three times with 1x wash buffer. MSD GOLD Read Buffer B (R60AM-2) was added to each well immediately prior to reading the plates on the MESO QuickPlex SQ 120MM machine. Raw signal values from the reference standard were fit to a 4-parameter logistic model with a 1/Y2 weighting, and antibody concentrations were determined by backfitting to the calibration curve and multiplying by the dilution factor. Reference control and sample fitting was performed through Meso Scale Discovery Workbench to generate concentrations in arbitrary units per mL (AU/mL). For SARS2 N IgG and WT S IgG, the concentration was then converted to World Health Organization arbitrary biological units per mL (BAU/mL) using the provided correction factors (0.00236 for N IgG and 0.00901 for WT S IgG). The LLOQ for the assay was defined using the lowest concentration of the reference standard for IgG with acceptable percent recovery (122% for N and 102% for WT S IgG) and adjusting for the 1:50,000 dilution used for most experimental samples. The resulting LLOQ in BAU/mL was 2.55 for N and 13.97 for WT S IgG. The LLOQ for 1:50,000 dilutions are shown on the longitudinal serology plots. A 1:5,000 dilution was used for a small subset of six plasma samples; all samples run at this dilution fell above the LLOQ calculated for 1:5,000 (0.26 BAU/mL for N and 1.40 BAU/mL for WT S IgG) and 1:50,000 dilutions. The negative control pre-pandemic plasma WT S IgG and N IgG concentrations fell below the LLOQ (N IgG geomean 0.114 and WT S IgG geomean 1.039 BAU/mL); the N IgG negative control is not shown (off scale for the y-axis minimum).

Validation of N IgG was performed by enzyme-linked immunosorbent assays (ELISA) (*8*). Plasma samples were heat-inactivated at 56°C for 30 minutes, aliquoted, and stored at 4°C prior to use. Corning 96-well half-area plates (ThermoFisher 3690) were coated with SARS2 N protein (AcroBiosystems NUN-C5221) at 1 μg/mL in phosphate-buffered saline (PBS) overnight at 4°C. The next day, the plates were blocked for 90 minutes at room temperature with 3% skim milk powder dissolved in 0.05% PBS-Tween 20 (ThermoFisher J20605-AP). After blocking, plasma samples were serially diluted two-fold across the plates in 1% skim milk powder dissolved in 0.05% PBS-Tween 20. Heat-inactivated, lab-generated pooled human plasma controls were also added and serially diluted across each plate. After a 90-minute incubation, the plates were washed 5x with 0.1% PBS-Tween 20 using a 405 Select Microplate Washer (BioTek 405TSUS). Secondary antibody (mouse IgG1 anti-human IgG Fc-HRP, Hybridoma Reagent Laboratory HP6043-HRP) was diluted 1:8,000 in 1% skim milk powder dissolved in 0.05% PBS-Tween 20 then added to each well and incubated for 60 minutes at room temperature. Plates were washed once more and developed using Ultra TMB-ELISA Substrate (ThermoFisher) for 5 minutes at room temperature. Reactions were quenched with 2N sulfuric acid ($H_2SO_4$). The optical density (OD) of the wells was then measured at 450 nm by a Spectramax M2 Plate Reader (Molecular Devices 89429-532) using SoftMax Pro software (v7.1.0).

Spectral flow cytometry surface immunophenotyping and analysis

Flow cytometry panels were developed and optimized for the detection of upper airway T cell subset markers and pMHC multimer staining using up to five different fluorophores for multimers. As noted above, enrollment was performed on a rolling basis, and not all subjects entered the study during the same calendar month. Some participants enrolled in the study prior to adoption of the final flow cytometry panels utilized for longitudinal PBMC and nasal swab staining. All nasal swab (MT and NP) samples were processed fresh, and flow cytometry was performed on the same day as sample collection. For cross-sectional analyses of bulk CD4 and CD8 T cells, all MT and NP swab samples and PBMC samples that met QC criteria and were stained with the final flow cytometry antibody panel were included. For longitudinal analyses of bulk CD4 or CD8 T cells, only samples from subjects who remained in the study for at least three months of sample collections were included.

Projected HLA allele population coverage

The Immune Epitope Database & Tools (IEDB) Population Coverage analysis resource (https://tools.iedb.org/population/) was used to calculate what proportion of the world population would be expected to respond to the pMHC multimers used in this study (*33*). All epitopes and MHC restricted alleles listed in Table S3 were submitted, 'World' was chosen for the population option, and 'Class I and II combined' was chosen for the calculation option.

Multimer (tetramer/pentamer) preparation and staining

The multimers used in this study were selected based on prior publications demonstrating their immunogenicity and specificity. The SARS2 pMHC-II LMI tetramer was the exception--it was demonstrated to be an immunodominant peptide by epitope mapping but had not previously been tested as a pMHC-II multimer (*41*). Additional considerations regarding multimer inclusion were the availability of reagents and the relative frequency of the HLA alleles of interest in our prospective study population. SARS2 CTF tetramer and biotinylated monomer were provided by the Jenkins' Lab at the University of Minnesota. SARS2 MEV, KTF, SPR, FTS, and TTD were obtained from ProImmune as pentamers conjugated to PE or APC. QLI, TDE, NLL were also obtained from ProImmune as tetramers conjugated to APC or PE. YLQ, KLW, and LMI were obtained from the NIH Tetramer Core Facility as monomers. KLW and LMI were requested as custom monomers using peptides were generated by Biopeptide Co., Inc. according to the NIH Tetramer Core requirements and based on the amino acid sequences listed in Tarke et al., 2021 (*41*). Influenza A GIL monomer was also obtained from the NIH Tetramer Core Facility. The Influenza A Class II PKY monomer was obtained from ProImmune. See Table S3 for additional details. All monomers provided by ProImmune and the NIH Tetramer Core were biotinylated, and aliquots were stored at -80°C until multimerized. Multimerization was performed based on the molecular weight and concentration of monomer provided. Streptavidin conjugated to the fluorophore of interest (BD, BioLegend) was added in excess (~4.5:1) and multimerization was performed at room temperature in the dark for 30 minutes. Phosphate buffered saline with 0.01% sodium azide was added to achieve a final working multimer concentration of 0.5 µM (for BUV737 and BB515 conjugates) or 1 µM (for PE, APC, BV421 conjugates). Working stocks were stored at 4°C.

Multimer validation

Multimer specificity was verified by multiple approaches and quantitative analyses. Negative controls tested included unstained samples, HLA-mismatched samples, and CLIP reagents from the Jenkins Lab and NIH Tetramer Core Facility (for class II responses). For the analyses shown, negative controls included multimer stained MT and NP samples from HLA-mismatched subjects and unstained PBMC samples. Single and dual (same multimer conjugated to two different fluorophores) multimer staining of MT, NP, and PBMC samples from HLA-matched subjects was also performed. For each multimer or allele (in the case of FTS/TTD and TDE/NLL), responses were compared between individual and concatenated data from HLA-matched samples versus negative control samples. Flow cytometry gates for MT and NP swab samples required that concatenated multimer responses from HLA-matched samples be at least three-times greater than concatenated responses from HLA-mismatched control samples. For PBMC, multimer gates were also set based on a similar threshold requirement; concatenated HLA-matched responses had to be at least three times greater than unstained control responses.

Spectral flow cytometry antigen-specific CD4 and CD8 memory T cell analysis

For longitudinal plotting of multimer-specific responses from nasal swab samples, 'Month 1' was relative to when multimer staining was initiated and denotes the first set of MT and NP swab samples stained with the specific multimer and using the final longitudinal flow cytometry panel, with subsequent time points being plotted as the time from 'Month 1'. Some participants who were enrolled in the study earlier provided nasal swab samples that were stained with an earlier version of the flow cytometry panel and/or fewer multimers as not all multimer reagents arrived at the same time. Given inherent variability in swab sampling, this earlier swab data was not included in longitudinal analyses. PBMC, however, were cryopreserved and run in batches after the final longitudinal flow cytometry panel was set and all multimer reagents had arrived. In cases where cryopreserved PBMC from the first study visit were collected before the longitudinal 'Month 1' for nasal swab samples, these PBMC samples were plotted as 'Month 0' on longitudinal PBMC plots, with subsequent PBMC samples plotted at the time (in months) from PBMC 'Month 0'. For longitudinal analyses of multimer-specific MT, NP, and PBMC CD4 or CD8 T cells, only samples from subjects who remained in the study for at least three months were included. Additional QC criteria were applied for inclusion in longitudinal group multimer response plots, as described above.

For the cross-sectional data shown in Fig. 1H-K and Fig. 2H-K, only data for HLA-typed individuals who provided paired MT, NP, and PBMC samples stained with the same multimers and antibody panel are shown. For correlation analyses (Fig. 1I-K, 2I-K, 4J), only samples that met quality control (QC) criteria were included.

Single cell high-dimensional flow cytometry analysis

Single-cell spectral flow cytometry data from 740 MT and NP samples (370 each) from 31 subjects were exported from OMIQ (Dotmatics) as csv files and were analyzed using Seurat (v5.0.3; (*42*)) in R (v4.3.2) following an approach adapted from Xu et al. (*43*). Raw fluorescence intensity values were asinh-transformed (cofactor 150) and used directly without additional normalization. Up to 5,000 cells per subject per anatomic site were randomly downsampled from a total of 11,453,283 CD4 (MT 453,546; NP 10,999,737) and 5,318,822 CD8 (MT 1,572,970; NP 3,745,852) T cells, yielding 285,228 CD4 and 307,121 CD8 T cells for Seurat object construction. Fourteen surface markers were used as variable features: CD69, CD103, CD49a, CXCR6, CCR6, CD161, CXCR3, CD127, CXCR5, CCR7, CD45RA, PD-1, CD25, and CD95. Data were scaled, followed by PCA (13 components), shared nearest neighbor graph construction (dims 1–13), Louvain clustering (resolution 0.2), and UMAP embedding (dims 1–13, default parameters), yielding 8 CD4 and 9 CD8 T cell clusters. For visualization, 50,000 cells were randomly subsampled from each object (Fig. 3D and Fig. 4F). Cluster identity was displayed using DimPlot split by anatomic site (MT and NP).

Cells binding more than one multimer other than those binding TDE/NLL or FTS/TTD, where binding to TDE vs NLL or FTS vs TTD could not be deconvoluted (same fluorophore used for each pair of multimers and same allele restriction), were excluded. SARS2 multimer-positive cells were then overlaid onto the UMAP embedding with multimer identity encoded by color. A downsampled overlay (Fig. 3E and Fig. 4G) included 2,236 CD4 and 848 CD8 pMHC-positive cells from HLA-matched subjects (n=25 CD4, n=23 CD8), plotted at increased point size relative to background cells.

Cluster composition was summarized as stacked bar plots (Fig. 3F and Fig. 4H) showing the percentage of cells from each cluster within four populations: bulk MT, multimer-positive MT, bulk NP, and multimer-positive NP. Bulk populations comprised all non-multimer-positive cells at each site. All CD4 (285,228) and CD8 (307,121) MT and NP T cells present in the respective Seurat objects were included in composition analyses without additional downsampling. Cell counts and cluster proportions are reported in the respective figures/legends.

$T_{RM}$ marker expression was visualized as dot plots (Fig. 3H and Fig. 4I) showing mean asinh fluorescence intensity (dot color, inverted plasma scale dark = high; cofactor 150, fixed color scale 0–6) and percentage of expressing cells (dot size). For bulk T cells, dot color was computed from the Seurat object background cells (277,900 CD4 and 300,807 CD8 bulk multimer-negative T cells), and dot size (% expressing) was computed as the percent of CD69 single positive or CD69 and other marker double-positive of parental CD69$^+$ non-naive memory T cells per sample from flow cytometry gate counts, averaged per anatomic site (MT or NP) across all samples. For multimer-positive T cells, dot color was computed from the full SARS2 multimer-positive T cell sets (23,318 pMHC-II positive CD4 and 15,279 pMHC- I positive CD8 cells), and dot size was computed as percent of CD69 single positive of CD69 and other marker double-positive of parental CD69$^+$ multimer-positive CD4/CD8 T cells per sample from OMIQ flow cytometry gate counts, averaged first across epitope specificities per subject per timepoint, then across subjects per anatomic site, requiring a minimum of three CD69$^+$ multimer-positive cells per sample. Bulk and multimer-positive dot plots were generated separately for CD4 and CD8 T cells, grouped by anatomic site.

Data Analysis and Statistics

Flow cytometry data acquisition was performed in SpectroFlo (Cytek, v2 or higher). Flow cytometry analysis was performed in FlowJo (BD, v10 or higher) and OMIQ software from Dotmatics (www.omiq.ai, www.dotmatics.com). Statistical analyses were performed in GraphPad Prism (v10 or higher) and R (v4.3.2 or higher).

### Supplementary Figure 1

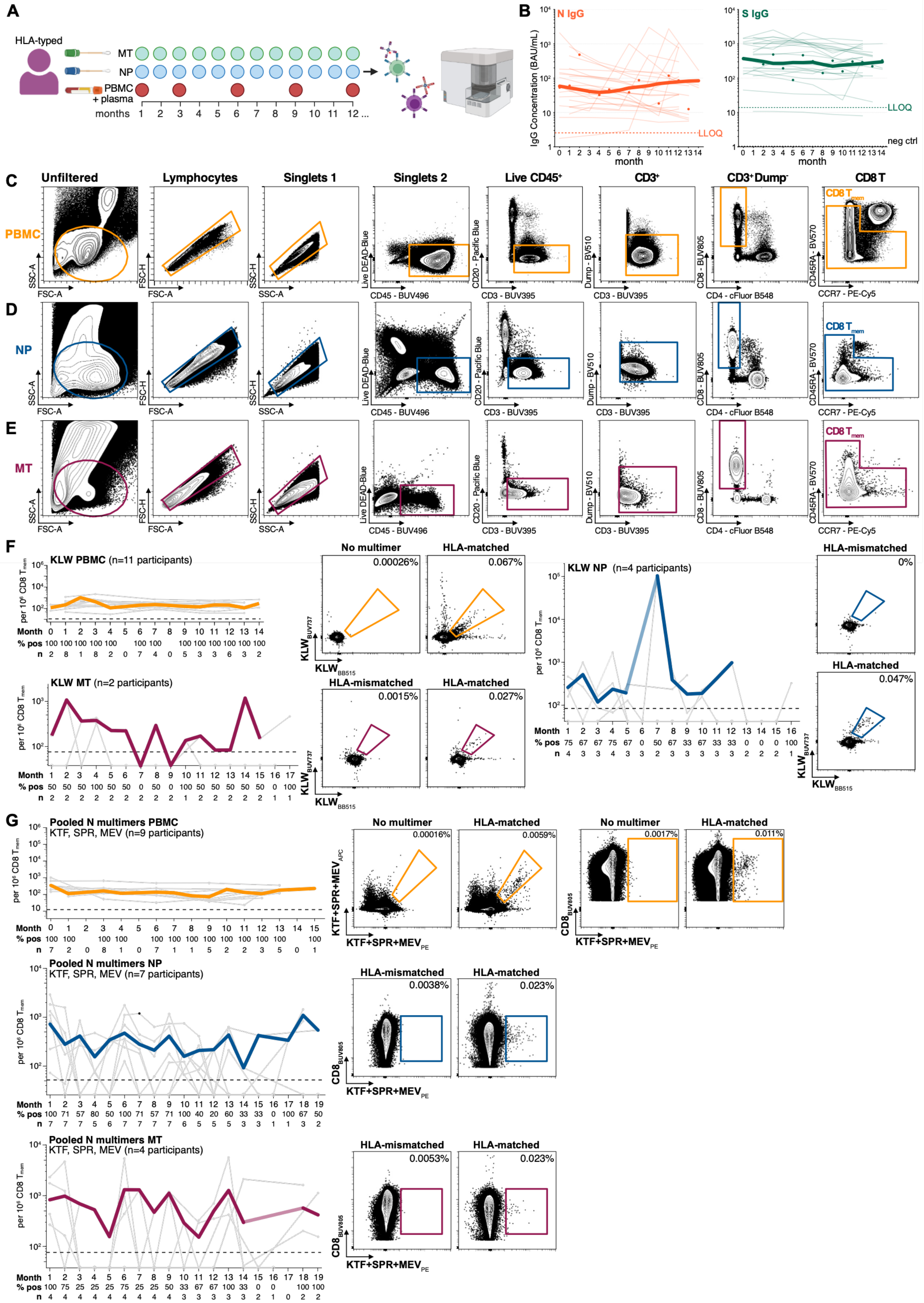

**Fig. S1. SARS2-specific CD8 T cell memory is durable across tissues in the absence of frequent infections.**

Related to Figure 1.

A. Diagram illustrating study sample types and collection time points. All study participants were HLA-typed. Plasma was utilized for serologic testing. MT and NP swabs, PBMC were used for pMHC-I and pMHC-II multimer staining. Spectral flow cytometry was the primary readout for multimer responses and was performed using a 5-laser Cytek Aurora spectral analyzer.

B. Group (n = 23) level longitudinal plasma SARS2 N and WT S IgG titers by MSD V-plex; values shown were converted to World Health Organization BAU/mL using the conversion factor provided by MSD. Thick line = LOWESS-smoothed geomean; circles = geomean values for each month that contributed to the smoothed geomean line. LLOQ = lower limit of quantification for the assay based on the lowest concentration of the standards provided with the kit. Neg ctrl (dotted black line) = IgG titer calculated from pooled pre-pandemic plasma samples (the value for N IgG falls below the y-axis minimum for the plot).

C-E. Flow cytometry gating strategies for memory CD8 T cells in (C) PBMC, (D) NP and (E) MT swab samples. The immediate downstream multimer gates are shown in the main text and supplementary figures.

F-G. Left: Longitudinal frequencies for SARS2 pMHC-I multimer-specific memory CD8 T cells in PBMC, NP and MT swabs of HLA-matched subjects and example flow cytometry plots for HLA-matched subjects versus HLA-mismatched negative controls for the (F) KLW and (G) nucleocapsid (KTF, SPR, MEV) multimers. In G, longitudinal nucleocapsid (N)-specific memory CD8 T cell frequencies for the KTF, SPR, and MEV pMHC-I multimers are plotted together as pooled N responses. Only subjects with at least 3 positive multimer responses (> or = 2 multimer-positive cells and > 100 CD8 T cells/MT swab or > 1,000 CD8 T cells/NP swab) were included in the longitudinal group plots; number of participants labeled on each plot. Thin gray lines = responses for individual subjects; solid gray dots = positive responses from PBMC or swabs with sufficient CD8 T cells, solid black dots = positive responses from swabs with insufficient CD8 T cells, open gray circles (floored) = negative responses from swabs with sufficient CD8 T cells. Thick colored line = group geomean (lighter colored line connects data points when group geomean calculations were not possible); only solid gray dots contributed to geomean calculations. Dashed black line = threshold for positivity; y-axis minimum (floor) = half this value. % pos = percent positive responders for each time point. Right: Example flow cytometry plots for SARS2 pMHC-I multimer-specific memory CD8 T cells in PBMC, NP and MT swabs of HLA-matched subjects or unstained (PBMC)/HLA-mismatched (NP, MT) controls for the (F) KLW and (G) KTF, SPR, and MEV multimers.

Supplementary Figure 2

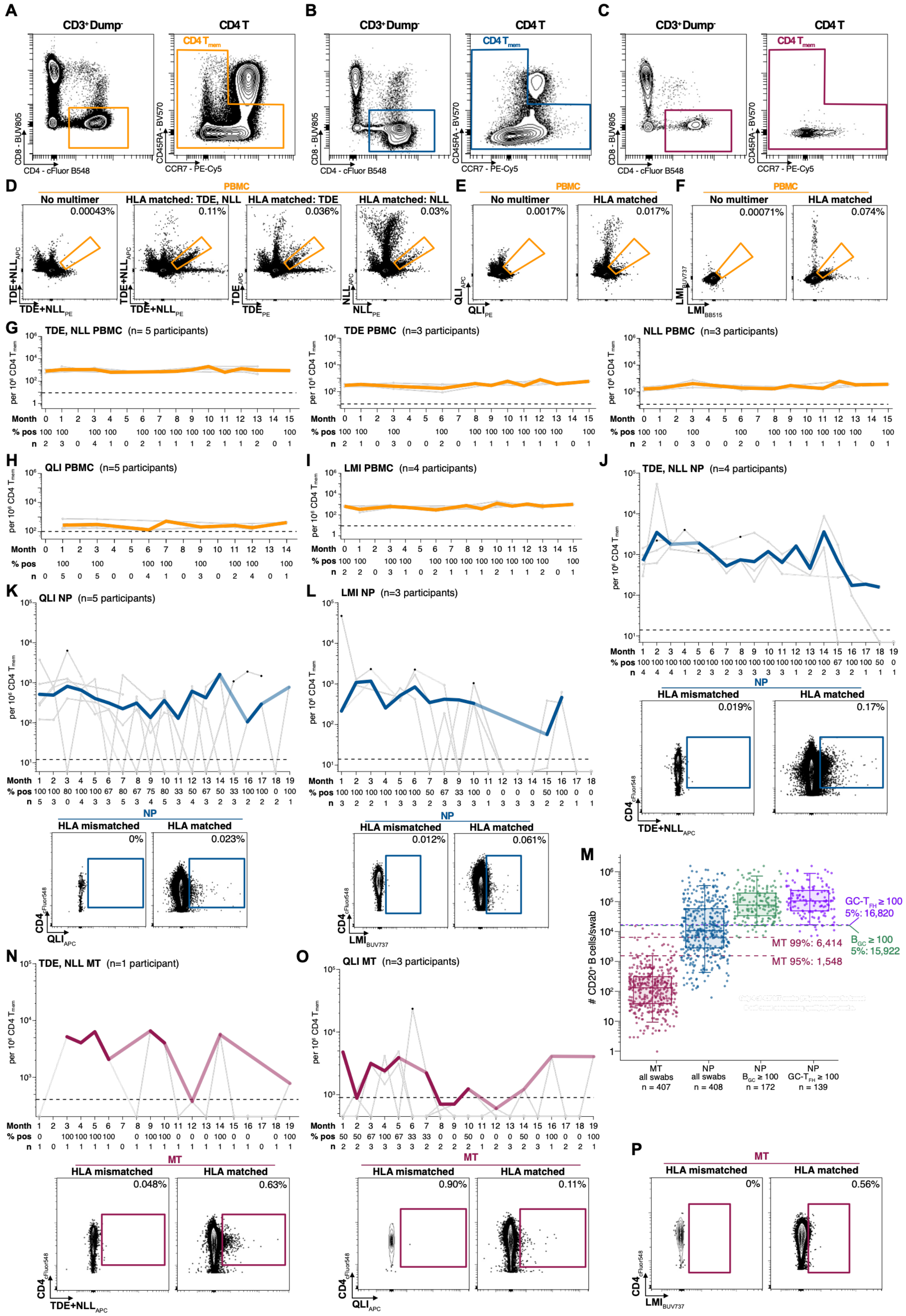

**Fig. S2. SARS2-specific CD4 T cell memory is durable across blood and upper airway tissues**

Related to Figure 2. A-C. Flow cytometry gating strategies for memory CD4 T cells in (A) PBMC, (B) NP and (C) MT swab samples. The upstream gates are the same as shown in Fig. S1C-E. The immediate downstream multimer gates are shown in the main text and supplementary figures.

D-F. Example concatenated flow cytometry plots for SARS2 pMHC-II multimer-specific memory CD4 T cells in dual stained PBMC of HLA-matched subjects or unstained PBMC from the same subjects for the (D) TDE, NLL (spike), (E) QLI (spike), and (F) LMI (non-spike) multimers.

G-I. Longitudinal frequencies of SARS2 pMHC-II multimer-specific memory CD4 T cells in PBMC of the same HLA-matched subjects as D-F for the (G) TDE, NLL, (H) QLI, and (I) LMI multimers. Thin gray lines = responses for individual subjects; solid gray dots = positive responses. Thick colored line = group geomean. Dashed black line = threshold for positivity; y-axis minimum (floor) = half this value. % pos = percent positive responders for each time point.

J-L. Top: Longitudinal frequencies of SARS2 pMHC-II multimer-specific memory CD4 T cells in NP swabs of HLA-matched subjects for the (J) TDE, NLL, (K) QLI, and (L) LMI multimers. Only subjects with at least 3 positive multimer responses (> or = 2 multimer-positive cells and > 1,000 CD4 T cells/NP swab) were included in the longitudinal group plots; number of participants labeled on each plot. Thin gray lines = responses for individual subjects; solid gray dots = positive responses from swabs with sufficient CD4 T cells, solid black dots = positive responses from swabs with insufficient CD4 T cells, open gray circles (floored) = negative responses from swabs with sufficient CD4 T cells. Thick colored line = group geomean (lighter colored line connects data points when group geomean calculations were not possible); only solid gray dots contributed to geomean calculations. Dashed black line = threshold for positivity; y-axis minimum (floor) = half this value. % pos = percent positive responders for each time point. Bottom: Example flow cytometry plots for concatenated NP swab data for HLA-matched subjects or HLA-mismatched controls for the (J) TDE, NLL, (K) QLI, and (L) LMI multimers.

M. A total B cell cutoff was used to classify NP swabs as adenoid vs non-adenoid NP samples. B cells per MT swab are shown in maroon. The 95th and 99th percentile cutoffs for B cells observed in MT swabs are shown as dashed maroon lines with the number of cells listed; MT 95%: 1,548 and MT 99% 6,414. NP swabs with at least 100 germinal center B cells ($B_{GC}$) or GC T follicular helper cells (GC-$T_{FH}$) are shown in green and purple, respectively. The 5th percentile cutoffs for total B cells in NP swabs with 100 or more $B_{GC}$ (15,922) and GC-$T_{FH}$ (16,820) are shown as labeled, dashed lines. Box, error bar, and whiskers = range, IQR, and median for each swab category; n = number of swabs per category. See Methods for additional details.

N-P. Top: Longitudinal frequencies of SARS2 pMHC-II multimer-specific memory CD4 T cells in MT swabs of HLA-matched subjects for the (N) TDE, NLL and (O) QLI multimers. Only subjects with at least 3 positive multimer responses (> or = 2 multimer-positive cells and > 100 CD4T cells/MT swab) were included in the longitudinal group plots; number of participants labeled on each plot. Thin gray lines = responses for individual subjects; solid gray dots = positive responses from swabs with sufficient CD4 T cells, solid black dots = positive responses from swabs with insufficient CD4 T cells, open gray circles (floored) = negative responses from swabs with sufficient CD4 T cells. Thick colored line = group geomean (lighter colored line connects data points when group geomean calculations were not possible); only solid gray dots contributed to geomean calculations. Dashed black line = threshold for positivity; y-axis minimum (floor) = half this value. % pos = percent positive responders for each time point. Bottom: Example flow cytometry plots for concatenated MT swab data for HLA-matched subjects or HLA-mismatched controls for the (N) TDE, NLL, (O) QLI, and (P) LMI multimers.

### Supplementary Figure 3

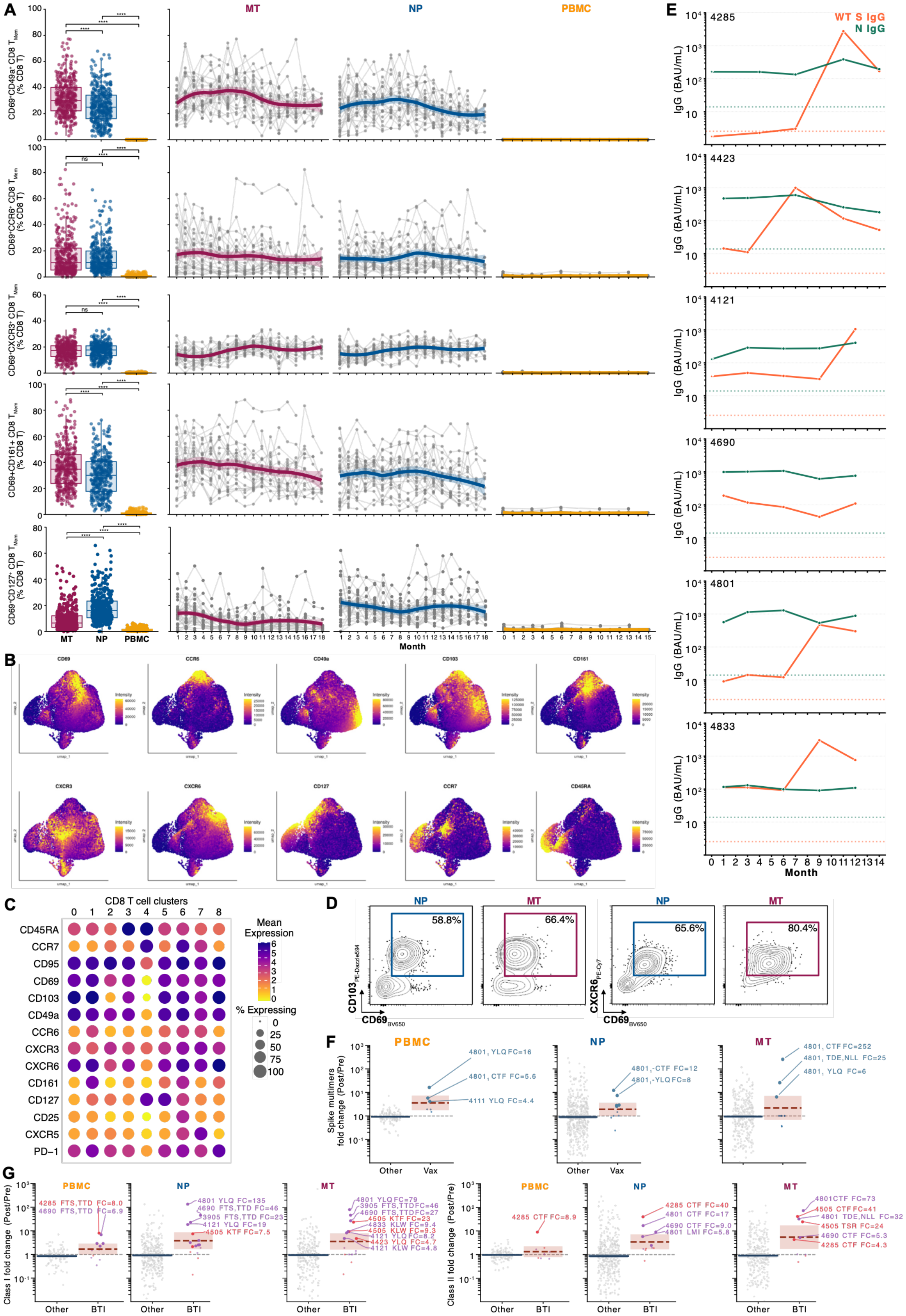

**Fig. S3. Upper airway CD8 T cell phenotypes are stable and SARS2-specific memory is boosted following Ag-exposure**

Related to Figure 3.

A. Longitudinal stability of the co-expression of CD69 and other surface markers by bulk memory CD8 T cells in PBMC, NP and MT swabs for the markers CD49a, CCR6, CXCR3, CD161, CD127.

B. Feature plots for total CD8 T cells related to Fig. 3D UMAPs.

C. Unscaled dot plot of marker expression for total CD8 T cells by Seurat cluster; related to Fig. 3D.

D. Example flow cytometry plots of $T_{RM}$ marker expression CD69 x CD103 and CD69 x CXCR6 by flu A GIL pMHC-I multimer-specific memory CD8 T cells in n = 70 NP and n = 70 MT swabs of HLA-matched subjects. % of parental (non-naive CD8 T cells) is shown.

E. SARS2 IgG plasma serology data for reported and suspected infection events. Plasma SARS2 N and WT S IgG titers by MSD V-plex; values shown were converted to WHO BAU/mL using the conversion factor provided by MSD. LLOQ = lower limit of quantification for the assay based on the lowest concentration of the standards provided with the kit dashed lines (orange for N IgG, green for S IgG).

F. Fold-change (FC) data for SARS2 spike pMHC-I and pMHC-II multimer responses related to reported vaccination events versus all other non-event time points split by site (PBMC, NP, MT). Ag exposure events with associated FC 2 or greater are shown as larger dots. Only Ag exposure events with associated FC 4 or greater are labeled. Geomean lines and confidence intervals are shown for each group. Dashed gray line at FC = 1. See Methods for additional details.

G. Fold-change data for SARS2 (spike and non-spike) pMHC-I and pMHC-II multimer responses related to reported (red) and suspected (purple) SARS2 infections versus all other non-event time points split by site (PBMC, NP, MT). Ag exposure events with associated FC 2 or greater are shown as larger dots. Only Ag exposure events with associated FC 4 or greater are labeled. Geomean lines and confidence intervals are shown for each group. Dashed gray line at FC = 1. See Methods for additional details.

### Supplementary Figure 4

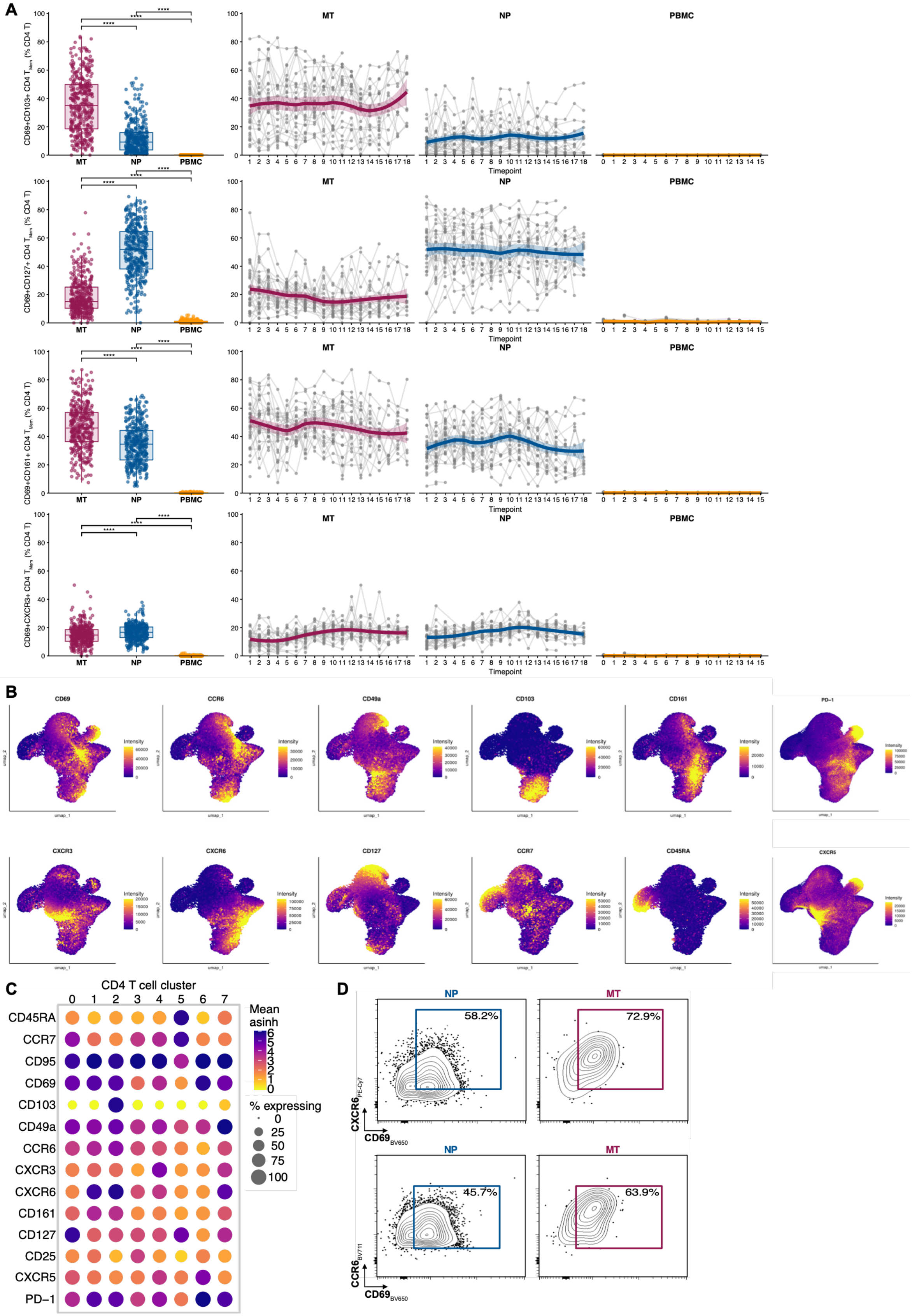

**Fig. S4. Upper airway CD4 T cells have stable phenotypes but display greater diversity across upper airway tissues**

Related to Figure 4.

A. Longitudinal stability of the expression of CD69 and other surface markers by bulk memory CD4 T cells in PBMC, NP and MT swabs for the markers CD103, CD127, CD161, CXCR3.

B. Feature plots for total CD4 T cells related to Fig. 4F UMAPs.

C. Unscaled dot plot of marker expression for total CD4 T cells by Seurat cluster; related to Fig. 4F.

D. Example flow cytometry plots of $T_{RM}$ marker expression CD69 x CXCR6 and CD69 x CCR6 by flu A PKY pMHC-II multimer-specific memory CD4 T cells in n = 47 NP and n = 47 MT swabs of HLA-matched subjects.

**Table S1.**

Study cohort demographics and relevant clinical history.

| | | |
|---|---|---|
| **Participant type** | Longitudinal, n (%) | 29 (85.3) |
| | Control, n (%) | 5 (14.7) |
| **Age (years)** | Median (IQR) | 37 (32.3 - 46.5) |
| **Sex** | Male, n (%) | 16 (47.1) |
| | Female, n (%) | 18 (52.9) |
| **Race** | White/Caucasian | 22 (64.7) |
| | Asian | 5 (14.7) |
| | More than one race | 4 (11.8) |
| | Deline to state | 3 (8.8) |
| **Ethnicity** | Hispanic | 9 (26.5) |
| | Non-Hispanic | 25 (73.5) |
| **Number of SARS2 vaccinations per participant at study entry** | Median (IQR) | 4 (3-4) |
| | Mean (Std Dev) | 3.6 (1.1) |
| **Number of SARS2 infections per participant at study entry** | Median (IQR) | 1 (1-2) |
| | Mean (Std Dev) | 1.3 (0.9) |
| **Participants who reported SARS2 infections during study period** | n (%) | 3 (10.3) |
| **Participants who reported COVID vaccines during study period** | n (%) | 6 (20.7) |
| **Participant who reported seasonal influenza vaccines during study period** | n (%) | 10 (34.5) |

Abbreviations: n, number of subjects or events; %, percentage; IQR, interquartile range; Std Dev, standard deviation. For reported infections and vaccinations during the study period, % calculation applies to longitudinal (non-control) participants.

**Table S2.**
Pertinent HLA allele information for the study cohort.

| Study ID | HLA-DPB1 Allele 1 | HLA-DPB1 Allele 2 | HLA-DRB1 Allele 1 | HLA-DRB1 Allele 2 | HLA-A Allele 1 | HLA-A Allele 2 | HLA-B Allele 1 | HLA-B Allele 2 |
|---|---|---|---|---|---|---|---|---|
| 0692 | DPB1*04:01 | DPB1*02:01 | DRB1*01:02 | DRB1*03:01 | A*01:01 | A*03:01 | B*07:02 | B*08:01 |
| 3905 | DPB1*02:01 | DPB1*15:01 | DRB1*04:08 | DRB1*11:04 | A*01:01 | A*01:01 | B*08:01 | B*35:02 |
| 4111 | DPB1*03:01 | DPB1*04:01 | DRB1*08:01 | DRB1*12:01 | A*02:01 | A*02:01 | B*44:02 | B*51:01 |
| 4121 | DPB1*02:01 | DPB1*03:01 | DRB1*07:01 | DRB1*13:02 | A*02:01 | A*66:01 | B*40:01 | B*50:01 |
| 4165 | DPB1*04:01 | DPB1*04:02 | DRB1*08:02 | DRB1*13:01 | A*02:01 | A*26:01 | B*35:12 | B*44:02 |
| 4169 | DPB1*03:01 | DPB1*04:01 | DRB1*03:01 | DRB1*04:01 | A*29:02 | A*31:01 | B*45:01 | B*51:01 |
| 4249 | DPB1*02:01 | DPB1*04:01 | DRB1*15:01 | DRB1*16:01 | A*02:01 | A*03:01 | B*44:02 | B*55:01 |
| 4284 | DPB1*03:01 | DPB1*04:02 | DRB1*03:01 | DRB1*15:01 | A*02:01 | A*24:02 | B*07:02 | B*50:01 |
| 4285 | DPB1*02:01 | DPB1*04:02 | DRB1*09:01 | DRB1*09:01 | A*01:01 | A*24:02 | B*15:11 | B*37:01 |
| 4311 | DPB1*02:01 | DPB1*02:01 | DRB1*01:01 | DRB1*13:02 | A*02:01 | A*31:01 | B*40:01 | B*44:12 |
| 4379 | DPB1*04:01 | DPB1*05:01 | DRB1*01:01 | DRB1*01:01 | A*25:01 | A*33:01 | B*56:01 | B*57:01 |
| 4423 | DPB1*04:01 | DPB1*04:01 | DRB1*04:01 | DRB1*11:02 | A*02:01 | A*24:02 | B*15:01 | B*39:01 |
| 4466 | DPB1*02:01 | DPB1*04:01 | DRB1*04:01 | DRB1*04:01 | A*02:01 | A*25:01 | B*40:01 | B*44:02 |
| 4484 | DPB1*04:02 | DPB1*05:01 | DRB1*01:01 | DRB1*07:01 | A*03:01 | A*03:01 | B*07:02 | B*35:01 |
| 4505 | DPB1*04:02 | DPB1*13:01 | DRB1*01:01 | DRB1*07:01 | A*02:01 | A*03:01 | B*35:01 | B*35:03 |
| 4549 | DPB1*04:02 | DPB1*11:01 | DRB1*07:01 | DRB1*15:03 | A*03:01 | A*03:01 | B*35:01 | B*44:03 |
| 4552 | DPB1*04:01 | DPB1*04:02 | DRB1*07:01 | DRB1*07:01 | A*01:01 | A*29:02 | B*44:03 | B*57:01 |
| 4562 | DPB1*04:01 | DPB1*04:02 | DRB1*04:11 | DRB1*14:06 | A*02:01 | A*03:01 | B*35:01 | B*52:01 |
| 4599 | DPB1*01:01 | DPB1*05:01 | DRB1*15:02 | DRB1*15:02 | A*34:01 | A*24:02 | B*27:06 | B*40:02 |
| 4571 | DPB1*03:01 | DPB1*04:01 | DRB1*01:03 | DRB1*15:01 | A*02:01 | A*11:01 | B*07:02 | B*38:01 |
| 4615 | DPB1*04:01 | DPB1*04:01 | DRB1*03:01 | DRB1*03:01 | A*33:03 | A*33:03 | B*58:01 | B*58:01 |
| 4648 | DPB1*02:02 | DPB1*05:01 | DRB1*01:01 | DRB1*08:03 | A*02:01 | A*02:06 | B*27:05 | B*54:01 |
| 4690 | DPB1*04:01 | DPB1*19:01 | DRB1*04:01 | DRB1*04:04 | A*01:01 | A*02:01 | B*41:01 | B*44:02 |
| 4714 | DPB1*05:01 | DPB1*13:01 | DRB1*08:03 | DRB1*08:03 | A*02:01 | A*02:07 | B*40:01 | B*46:01 |
| 4726 | DPB1*04:01 | DPB1*04:02 | DRB1*13:01 | DRB1*14:06 | A*02:01 | A*26:01 | B*38:01 | B*51:01 |
| 4795 | DPB1*04:01 | DPB1*14:01 | DRB1*11:01 | DRB1*15:01 | A*03:01 | A*23:01 | B*07:02 | B*49:01 |
| 4801 | DPB1*01:01 | DPB1*04:01 | DRB1*03:01 | DRB1*15:01 | A*01:01 | A*02:01 | B*08:01 | B*51:01 |
| 4818 | DPB1*04:01 | DPB1*10:01 | DRB1*01:01 | DRB1*09:01 | A*02:01 | A*02:01 | B*40:01 | B*51:01 |
| 4833 | DPB1*02:01 | DPB1*02:01 | DRB1*04:01 | DRB1*11:01 | A*02:01 | A*29:02 | B*44:02 | B*44:02 |
| 4913 | DPB1*04:01 | DPB1*03:01 | DRB1*04:06 | DRB1*09:01 | A*02:01 | A*24:02 | B*39:05 | B*50:02 |
| 4951 | DPB1*04:01 | DPB1*04:01 | DRB1*01:01 | DRB1*04:01 | A*03:01 | A*03:01 | B*07:02 | B*57:01 |
| 4981 | DPB1*04:01 | DPB1*04:02 | DRB1*01:01 | DRB1*04:04 | A*02:20 | A*24:02 | B*35:01 | B*40:02 |
| 5555 | DPB1*04:01 | DPB1*04:01 | DRB1*15:01 | DRB1*15:01 | A*02:01 | A*24:02 | B*35:02 | B*44:02 |
| 7373 | DPB1*02:01 | DPB1*02:01 | DRB1*01:01 | DRB1*04:05 | A*02:05 | A*24:02 | B*38:01 | B*58:01 |

**Table S3.**

SARS-CoV-2 (SARS2) and influenza A pMHC-I and pMHC-II multimers used in the study.

| Class II Allele | Antigen | Peptide sequence | Fluor 1 | Fluor 2 | Virus | Source | Format | Product info |
|---|---|---|---|---|---|---|---|---|
| DPB1*04:01/ 04:02 | $S_{166-182}$ | CTFEYVSQPFLMDLE | APC | BV421 | SARS2 | Jenkins lab | monomer, tetramer | N/A |
| DRB1*01:01 | $M_{172-188}$ | TSRTLSYYKLGASQRVA | APC | PE | SARS2 | ProImmune | tetramer | ProT2® MHC Class II Tetramers peptide code 4407 |
| DRB1*04:01 | $S_{993-1010}$ | QLIRAAEIRASANLAATK | APC | PE | SARS2 | ProImmune | tetramer | ProT2® MHC Class II Tetramers peptide code 4331A |
| DRB1*15:01 | $S_{866-880}$ | TDEMIAQYTSALLAG | APC | PE | SARS2 | ProImmune | tetramer | ProT2® MHC Class II Tetramers peptide code 4406 |
| DRB1*15:01 | $S_{751-767}$ | NLLLQYGSFCTQLNRAL | APC | PE | SARS2 | ProImmune | tetramer | ProT2® MHC Class II Tetramers peptide code 4414 |
| DRB1*15:01 | nsp12 | LMIERFVSLAIDAYP | BUV737 | BB515 | SARS2 | NIH Tetramer Core | tetramer | custom biotinylated monomer |
| DRB1*07:01 | $HA_{307-319}$ | PKYVKQNTLKLAT | BV421 | BUV737 | Influenza A | ProImmune | tetramer | ProM2® MHC Class II Monomers, biotinylated peptide code 357C |

| Class I Allele | Antigen | Peptide sequence | Fluor 1 | Fluor 2 | Virus | Source | Format | Product Info |
|---|---|---|---|---|---|---|---|---|
| A*01:01 | $Orf3_{207-215}$ | FTSDYYQLY | PE | APC | SARS2 | ProImmune | pentamer | Catalog Pro5® MHC Class I Pentamers peptide code 4355 |
| A*01:01 | $Orf1ab_{1637-1646}$ (nsp3) | TTDPSFLGRY | PE | APC | SARS2 | ProImmune | pentamer | Catalog Pro5® MHC Class I Pentamers peptide code 4381 |
| A*02:01 | $S_{269-277}$ | YLQPRTFLL | PE | BB515 | SARS2 | NIH Tetramer Core | premade monomer | premade class I biotinylated monomer |
| A*02:01 | $ORF1ab_{3886-3894}$ (nsp7) | KLWAQCVQL | BUV737 | BB515 | SARS2 | NIH Tetramer Core | custom monomer | custom biotinylated monomer |
| A*03:01 | $N_{362-370}$ | KTFPPTEPK | PE | APC | SARS2 | ProImmune | pentamer | Catalog Pro5® MHC Class I Pentamers peptide code 4356A |
| B*07:02 | $N_{105-113}$ | SPRWYFYYL | PE | APC | SARS2 | ProImmune | pentamer | Catalog Pro5® MHC Class I Pentamers peptide code 4351 |
| B*40:01 | $N_{322-331}$ | MEVTPSGTWL | PE | APC | SARS2 | ProImmune | pentamer | Catalog Pro5® MHC Class I Pentamers peptide code 4328 |
| A*02:01 | $M1_{58-66}$ | GILGFVFTL | BB515 | BUV737 | Influenza A | NIH Tetramer Core | premade monomer | premade class I biotinylated monomer |

**Table S4.**

Flow cytometry antibodies used in the study for all MT and NP swab samples collected before January 20, 2026.

**T Cell Panel B version 2**

| Channel | Target | Fluorochrome | Peak | Clone | Vendor, Cat# | Dilution |
|---|---|---|---|---|---|---|
| UV | CD3 | BUV395 | UV2 | UCHT1 | BD, 563546 | 1:500 |
| | VIABILITY | Live-Dead BLUE | UV6 | | Thermo, L23105 | 1:1,000 |
| | CD45 | BUV496 | UV7 | HI30 | BD, 569101 | 1:500 |
| | CD49a | BUV563 | UV9 | TS2/7 | BD, 755216 | 1:200 |
| | | | UV10 | | | |
| | Multimer 1 | BUV737 | UV14 | | | See Methods |
| | CD8 | BUV805 | UV16 | SK1 | BD, 612889 | 1:500 |
| Violet | Multimer 2 | BV421 | V1 | | | See Methods |
| | CD20 | Pacific Blue | V3 | 2H7 | BioLegend, 302320 | 1:200 |
| | Dump | BV510 | V7 | CD14 (63D3)<br>CD16 (3G8)<br>CD56 (5.1H11) | BioLegend, 367124<br>BioLegend, 302048<br>BioLegend, 362534 | 1:500 each |
| | CD45RA | BV570 | V8 | HI100 | BioLegend, 304132 | 1:500 |
| | CXCR3 | BV605 | V10 | G025H7 | BioLegend, 353728 | 1:200 |
| | CD69 | BV650 | V11 | FN50 | BioLegend, 310934 | 1:200 |
| | CCR6 | BV711 | V13 | G034E3 | BioLegend, 353436 | 1:200 |
| | PD-1 | BV785 | V15 | EH12.2H7 | BioLegend, 329930 | 1:200 |
| Blue | Multimer 3 | BB515 | B1 | | | See Methods |
| | CD4 | cFluor 548 | B3 | SK3 | Cytek, R27-20043 | 1:500 |
| | CD95 | RB613 | B6 | DX2 | BD, 758755 | 1:200 |
| | CD25 | BB700 | B9 | BC96 | BD, 567481 | 1:200 |
| | | | B14 | | | |
| Yellow/ Green | Multimer 4 | PE | YG1 | | | See Methods |
| | CD103 | PE-Dazzle594 | YG3 | Ber-ACT8 | BioLegend, 350224 | 1:200 |
| | CCR7 | PE-Cy5 | YG5 | G043H7 | BioLegend, 353272 | 1:200 |
| | CD127 | PE-Fire 700 | YG7 | A019D5 | BioLegend, 351366 | 1:200 |
| | CXCR6 | PE-Cy7 | YG9 | K041E5 | BioLegend, 356012 | 1:200 |
| Red | Multimer 5 | APC | R1 | | | See Methods |
| | CD161 | R718 | R4 | DX12 | BD, 751652 | 1:200 |
| | CXCR5 | APC-F750 | R7 | J252D4 | BioLegend, 356946 | 1:200 |
| | CD38 | APC-F810 | R8 | HB-7 | BioLegend, 356644 | 1:200 |

**Table S5.**

Flow cytometry antibodies used in the study for all PBMC samples, and MT and NP swab samples collected January 20, 2026 or later.

**T Cell Panel B version 3**

| Channel | Target | Fluorochrome | Peak | Clone | Vendor, Cat# | vol (uL) per mL |
|---|---|---|---|---|---|---|
| **UV** | CD3 | BUV395 | UV2 | UCHT1 | BD, 563546 | 1:500 |
| | VIABILITY | Live-Dead BLUE | UV6 | | Thermo, L23105 | 1:1,000 |
| | CD45 | BUV496 | UV7 | HI30 | BD, 569101 | 1:500 |
| | CD49a | BUV563 | UV9 | TS2/7 | BD, 755216 | 1:200 |
| | **HLA-DR** | **BUV615** | **UV10** | **G46-6** | **BD, 751142** | **1:500** |
| | Multimer 1 | BUV737 | UV14 | | | See Methods |
| | CD8 | BUV805 | UV16 | SK1 | BD, 612889 | 1:500 |
| **Violet** | Multimer 2 | BV421 | V1 | | | See Methods |
| | CD20 | Pacific Blue | V3 | 2H7 | BioLegend, 302320 | 1:200 |
| | Dump | BV510 | V7 | CD14 (63D3)<br>CD16 (3G8)<br>CD56 (5.1H11) | BioLegend, 367124<br>BioLegend, 302048<br>BioLegend, 362534 | 1:500 each |
| | CD45RA | BV570 | V8 | HI100 | BioLegend, 304132 | 1:500 |
| | CXCR3 | BV605 | V10 | G025H7 | BioLegend, 353728 | 1:200 |
| | CD69 | BV650 | V11 | FN50 | BioLegend, 310934 | 1:200 |
| | CCR6 | BV711 | V13 | G034E3 | BioLegend, 353436 | 1:200 |
| | PD-1 | BV785 | V15 | EH12.2H7 | BioLegend, 329930 | 1:200 |
| **Blue** | Multimer 3 | BB515 | B1 | | | See Methods |
| | CD4 | cFluor 548 | B3 | SK3 | Cytek, R27-20043 | 1:500 |
| | CD95 | RB613 | B6 | DX2 | BD, 758755 | 1:200 |
| | CD25 | BB700 | B9 | BC96 | BD, 567481 | 1:200 |
| | **CD45RO** | **PerCP/Fire 806** | **B14** | **UCHL1** | **BioLegend, 304276** | **1:500** |
| **Yellow/ Green** | Multimer 4 | PE | YG1 | | | See Methods |
| | CD103 | PE-Dazzle594 | YG3 | Ber-ACT8 | BioLegend, 350224 | 1:200 |
| | CCR7 | PE-Cy5 | YG5 | G043H7 | BioLegend, 353272 | 1:200 |
| | CD127 | PE-Fire 700 | YG7 | A019D5 | BioLegend, 351366 | 1:200 |
| | CXCR6 | PE-Cy7 | YG9 | K041E5 | BioLegend, 356012 | 1:200 |
| **Red** | Multimer 5 | APC | R1 | | | See Methods |
| | CD161 | R718 | R4 | DX12 | BD, 751652 | 1:200 |
| | CXCR5 | APC-F750 | R7 | J252D4 | BioLegend, 356946 | 1:200 |
| | CD38 | APC-F810 | R8 | HB-7 | BioLegend, 356644 | 1:200 |

**Data S1. (separate file)**
This file contains the tabulated data behind the main text and supplementary figures other than high-dimensional flow cytometry single cell analyses.

**Data S2. (separate file)**
This file contains the tabulated data related to high-dimensional flow cytometry single cell analyses.
